# Sex-biased gene expression under sexually antagonistic and sex-limited selection

**DOI:** 10.1101/2024.11.29.622017

**Authors:** R. Axel W. Wiberg, Martyna K. Zwoinska, Philipp Kaufman, James M. Howie, Elina Immonen

## Abstract

Sex differences in gene expression are ubiquitous, evolve quickly, and are expected to underlie phenotypic sexual dimorphism. Despite long-standing interest, the impact of sex-specific selection on the transcriptome remains poorly understood. Here, we test fundamental questions on the role of constraints on gene expression evolution arising from the mode of selection and genetic architecture. We also test the relationship between sex-biased expression and evolved sexual dimorphism (SD). We assess these using body size selection lines in the seed beetle, *Callosobruchus maculatus,* that have evolved variation in SD in response to either sex-limited (SL) or sexually antagonistic (SA). We find that sex differences in the phenotypic responses and expression changes are generally well aligned. SL selection, despite a phenotypic response similar to SA selection in males, but not in females, resulted in a more extensive expression differentiation and increase of sex-biased expression than SA selection. These patterns show that SA selection imposes a transcriptomic constraint and is not required for sex-bias to evolve. Sex-biased transcripts show lower cross-sex correlations in expression changes than unbiased transcripts, suggesting greater sex differences in their underlying genetic architecture. Although male-biased transcripts are disproportionately affected when selection targeted males, we find no support for a transcriptome-wide association between sex-bias and SD. In the light of these unique experimental insights into how sex-specific selection on size changes adult transcription, our findings have important implications for inferring selection history and mode from patterns of sex-biased gene expression in natural populations.

## Introduction

In most species with separate sexes, female and male fitness is optimized through different strategies, causing selection for phenotypic sex differences. The resulting sexually dimorphic phenotypes are some of the most extraordinary in nature (Fairbairn 2007), including sexual display traits and weaponry (Emlen 2008), but also many physiological and life history traits expressed by both sexes, such as body size (Wedell et al. 2006; Austad & Fischer 2016; Hämäläinen et al. 2018). Evolution of sexual dimorphism (SD) depends on the interplay between selection favouring sex differences and the constraint imposed by the shared genetic architecture of the trait (Lande 1980; Fairbairn 2007; Pennell & Morrow 2013). An important mechanism that should facilitate the evolution of SD is sex-biased or sex-limited expression of the underlying genes (Ellegren & Parsch 2007; Parsch & Ellegren 2013; Grath & Parsch 2016). Sex-limited (SL, i.e. selection only on one sex, with neutral evolution of the other) selection is expected to limit sexual dimorphism at both phenotypic and molecular levels due to correlated evolutionary response of the opposite sex. In contrast, sexually antagonistic (SA; i.e. opposing directions of selection in the two sexes) selection is expected to be a chief selective force behind sex-biased gene expression because it generates a genomic conflict between the sexes over shared genes that differential gene regulation can resolve (Ellegren & Parsch 2007; Parsch & Ellegren 2013; Grath & Parsch 2016). Many studies have therefore sought a link between overall levels of sexual dimorphism, sex-biased gene expression and either ongoing or historical SA selection. However, there have been few experimental tests of these commonly held expectations, and interpreting process from patterns observed in nature is far from trivial. Moreover, the genetic architecture of gene regulation itself can constrain the evolution of sex-biased gene expression, but the scope and consequence of such constraints on modulating gene expression evolution are not well understood (Ayroles et al. 2009; Griffin et al. 2013; Cheng & Kirkpatrick 2016; Allen et al. 2018; Houle & Cheng 2021).

While some studies have found a positive association between sexual dimorphism and sex-biased expression (Pointer et al. 2013; Harrison et al. 2015; Toubiana et al. 2021), others have not (Scharmann et al. 2021; Khila et al. 2012). The link between SD and sex-biased expression is thus not straightforward, making it difficult to infer optimal expression levels for either sex (Immonen et al. 2017; Parker et al. 2019; Huylmans et al. 2021). Ruzicka et al. (2019) measured SA fitness variation directly, and showed that SA genes (i.e. genes harbouring SNPs with sexually antagonistic fitness effects) in *Drosophila melanogaster* are in fact *less* often sex-biased in expression than expected by chance, suggesting that sex-biased expression may not be a reliable indicator of sexually antagonistic loci. Meanwhile, in experimental evolution settings, sex-biased gene expression evolves in response to altered mating systems as predicted. However, results are often contradictory, with monogamy (and thus relaxed sexual conflict) resulting in the “feminisation” of gene expression in some cases (i.e. increased expression of female-biased genes, and reduced expression of male-biased genes; Hollis et al. 2014), or “masculinisation” in others (Immonen et al. 2014; Veltsos et al. 2017). The evidence for a consistent effect of a history of SA selection on sex-biased genes is similarly mixed. Studies of flycatchers (Dutoit et al. 2018) and seed beetles (Sayadi et al. 2019) have found excess of non-synonymous polymorphisms in sex-biased genes. This is in line with maintenance of genetic variation due to overall balancing selection from opposing selection pressures on males and females, a process expected from theory (Connallon & Clark 2012). Such patterns are consistent with a history of persistent sexual conflict, but also with any other form of antagonistic pleiotropy as well as relaxed selection (Dutoit et al. 2018; Dapper and Wade 2020).

From theory, it is not immediately clear what predictions should be. It is well established that phenotypic sexual dimorphism can change in response to sex-limited directional selection, or even sexually concordant selection, as long as there are quantitative asymmetries in genetic variation between the sexes available to selection, potentially making SA selection unnecessary (Fisher 1930; Lande 1980; Bonduriansky & Chenoweth 2009; Gosden et al. 2012; Cheng & Houle 2020; Houle & Cheng 2021; Kaufmann et al. 2021). In addition, the variable patterns observed across studies could partly be explained by differences in the evolutionary timescale used to assess the associations between sex-bias, sexual dimorphism and SA selection. If there is a time-lag between the evolution of sexual dimorphism and the resolution of sexual conflict over such dimorphism through sex-biased gene expression, then comparing very recently diverged populations or species may not yield the expected positive relationship. Finally, in comparative studies of sex-bias, it is typically not known what phenotypes selection has acted on, nor the exact form of selection, which are instead inferred from the data or assumed (Ellegren & Parsch, 2007; Grath & Parsch 2016; Mank 2017). It is therefore not clear what kind of signature SA and other forms of selection leave on gene expression, and the link with phenotypic sexual dimorphism remains poorly understood.

Here, we experimentally address how gene expression is altered in each sex in response to different modes of sex-specific selection on body size in the seed beetle *Callosobruchus maculatus*, by combining RNA-seq with replicated artificial selection (Kaufmann et al. 2021). Body size is a key life history trait and commonly sexually dimorphic across taxa (Fairbairn 2007; Wedell et al. 2006; Immonen et al. 2018). In *C. maculatus* it is differentially associated with fitness in the sexes: fecundity selection favours larger females and sexual selection typically favours smaller and more active males (Arnqvist & Tuda 2010; Berger et al. 2014; 2016). Most of the autosomal additive genetic variance for body size is shared between the sexes both in the ancestral study population of *C. maculatus* and in the selected lines (autosomal *r_MF_*> 0.92, Kaufmann et al. 2021; Kaufmann et al. 2023a). Sex-specific evolution of body size in this system therefore offers an excellent model to address how gene expression changes in the sexes in association with polygenic adaptation in the presence of this high cross-sex genetic correlation.

Full details of the artificial selection lines are given elsewhere (Kaufmann et al. 2021). Briefly, the modes of selection in our experiment include male-limited (towards smaller males: SL_m_↓, and larger males: SL_m_↑), female-limited (towards larger females: SL_f_↑) and SA selection (in the naturally observed direction of larger females and smaller males). We also include a control line (C) subjected to random selection but with otherwise similar demographic changes. The C line has largely maintained genetic variation for body size through the course of selection (Kaufmann et al. 2023a). A total of 10 generations of selection has produced a gradient of sexual size dimorphism (figure 1). SD changes range from 50% and 30% increase under SA and SL_m_↓ selection on smaller males, respectively, through no change under SL_f_↑ selection (due to a correlated increase in size in both sexes), to some 30% decrease under SL_m_↑ selection (due to a greater increase in male size compared to female size; Kaufmann et al. 2021; see figure S1). Y-linked variance accounted for changes in SD during the first 2-3 generations (Kaufmann et al. 2021, 2023a). While most efficiently increasing sexual size dimorphism, SA selection maintained significantly more autosomal genetic variance (additive and dominance) compared to male-limited selection for similar male but not female size (Kaufmann et al. 2023a). These findings corroborate the theoretical expectations that SA selection can both target loci with mutually beneficial effects on both sexes (by increasing SD) but also impose strong genomic constraints on shared genetic variation.

**Figure 1.**
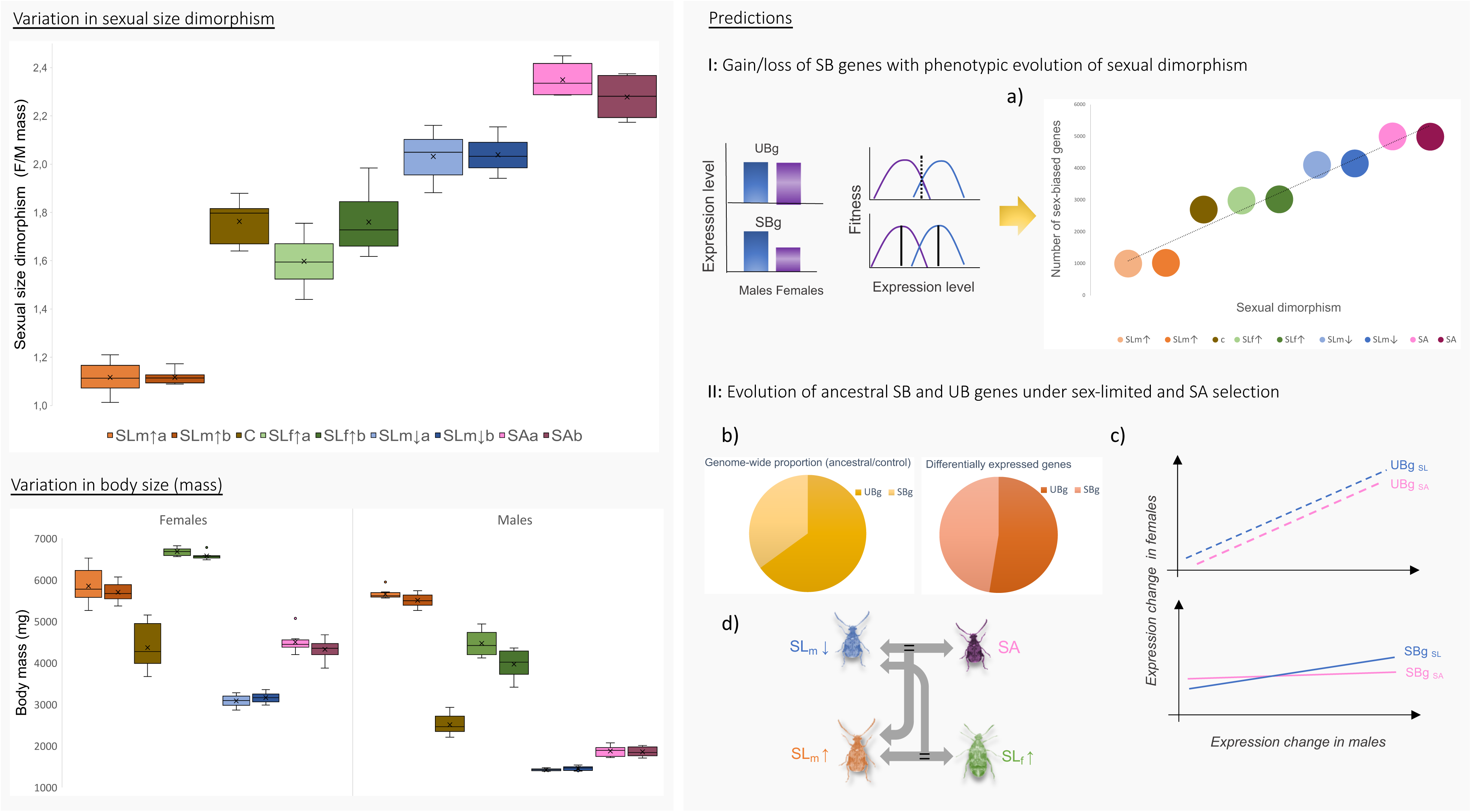
*Left column:* Variation in sexual size dimorphism (SSD) across the selection lines sequenced in this study (top), and the body mass distributions of males and females across the same lines (bottom). *Right column:* schematic illustrations of the predictions **A)** of a positive association between the number/proportion of sex-biased (SB) genes and phenotypic sexual dimorphism, **B)** that ancestrally sex-biased genes are disproportionately affected by sex-limited and SA selection and **C)** show lower cross-sex correlations in expression evolution compared to unbiased (UB) genes. SA selection should lead to fewer, but more sex-specific expression changes than sex-limit (SL) selection. **D)** Phenotypic and expression changes are associated in terms of the identity of transcripts.

Here, we take advantage of this known selection history, controlled demography and a range of phenotypic sexual dimorphism produced by selection (figure 1), to address the following questions: *Does the degree of sexual size dimorphism covary with transcriptome-wide sex-bias? How does the mode of selection affect the extent and parallelism of expression change, sex-biased gene expression, and cross-sex correlations in gene expression?*

We predict a positive association between the number/proportion of sex-biased transcripts and phenotypic sexual size dimorphism, under the premise that sex-biased expression is more readily gained when sexual dimorphism increases, and concomitantly lost when it decreases (figure 1A). We further expect that ancestrally sex-biased transcripts are disproportionately affected by sex-specific selection and should also show less correlated changes between the sexes compared to unbiased transcripts, under the expectation that they have more independent regulatory architectures between the sexes (figure 1B-C). SA selection should lead to overall fewer and less correlated expression changes between the sexes compared to sex-limited directional selection where, unlike under SA selection, both direct and correlated evolution has changed the body size of both sexes (Kaufmann et al. 2021). We also expect some consistency between the phenotypic response to selection and expression, both in terms of extent of changes and the identity of transcripts (figure 1D). For example, the SA and SLm↓ lines have undergone similar phenotypic changes in males, and we therefore expect that not only the replicate lines but even these different selection modes show an overlap of expression changes. Consistency in expression responses can however be limited by genetic redundancy predicted for polygenic traits that reduce parallel evolutionary responses in populations subject to similar selection pressures (Barghi et al. 2020). Also, similar selection but acting on a different sex is expected to reduce the consistency of expression changes if the sexes differ in the underlying genetic variances (Houle & Cheng 2021).

## Results

### Is there an association between sexual size dimorphism and transcriptome-wide sex-bias?

We evaluated expression differences between the sexes within each of the nine lines, in a total of 19,373 transcripts. Across all the lines, there is variation in the number of transcripts with significant sex-bias but no evidence for a transcriptome-wide association between the number or proportion of sex-biased transcripts and degree of sexual size dimorphism (SSD), for either male-biased (autosomes: rho = 0.21, p = 0.59, X-chromosome: rho = 0.13, p = 0.75) or female-biased transcripts (autosomes: rho = 0.16, p = 0.68, X- chromosome: rho = -0.18, p = 0.64; figure 2). Autosomal and X-linked transcript numbers show a similar rank order across the lines (figure 2). There was some evidence for a subtle, but significant, negative relationship between the transcriptome-wide magnitude of sex-bias (the median log of fold change in expression, logFC, between males and females) and SSD across all the lines for male-biased transcripts (rho = -0.74, p = 0.02) but no relationship in the female-biased transcripts (rho = -0.03, p = 0.95; figure S3).

**Figure 2.**
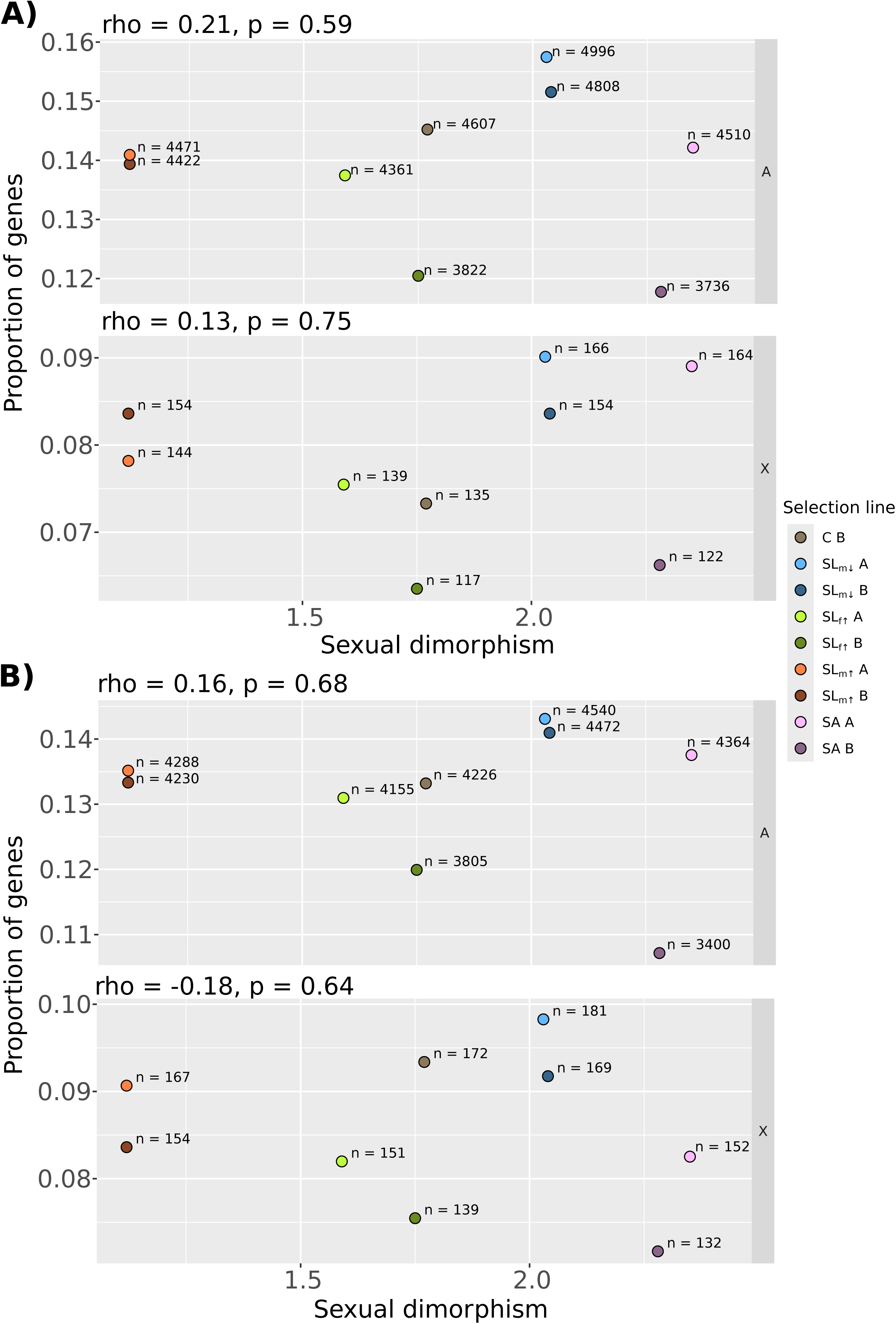
The proportion of **A)** male-biased and **B)** female-biased transcripts as a function of sexual dimorphism across selection line treatments. Inset text above each panel gives the result of a Spearman rank correlation test for the association between the proportion of transcripts and sexual dimorphism. Inset text next to each point give the absolute number of transcripts in each case. The data are split by chromosome category into autosomes – ‘A’, the X-chromsome – ‘X’.

### Turnover in sex-biased gene expression

The vast majority of transcripts show significant sex-bias in the same transcripts and in the same direction in all selection lines and the control (C) line (figure S2), as expected. Pearson’s correlation coefficients of the female-male logFC between each selection line and the C line are all >0.9 (all p < 0.001; figure S1). Interestingly, the overlap of sex-bias status is proportionally greater for female-biased transcripts, of which 2,466 (56% of those female- biased in C) transcripts are shared across all selection lines, compared to 1,883 male-biased transcripts (39% of those male-biased in C; figure S2), indicating a greater turnover of male- bias. Gains of novel sex-bias are most frequently line-specific and have occurred in thousands of transcripts across the lines (1,677 and 1,179 line-specific gains of male- and female-bias, respectively; figure S2). Consistent gains of novel sex-bias in both replicate lines of a given selection mode (i.e. in transcripts that remain unbiased in all other lines) are relatively few but more frequent under male-limited selection modes compared to the selection modes acting also or exclusively on females (gain of sex-bias in 109, 95, 36 and 0 transcripts in the SL_m_↓ A/B, SL_m_↑ A/B, SA A/B and SL_f_↑ A/B, respectively; figure S2). There is however no evidence for sex-by-line interaction effects at these loci (at FDR < 0.05) between the C line and both selection line replicates of these modes, apart from one transcript (in the C vs SL _m_↓ A as well as C vs SL_m_↓ B). Evidence for interaction effects was generally weak, and found in individual selection line replicates in only a handful of cases (up to 7 transcripts in the C vs SL_m_↓ A). In line with this, also reversal of sex-bias is rare, with consistent evidence in only two transcripts across both selection line replicates (one is male-biased in C and female- biased in the SL_m_↑, another is female-biased in C and male-biased in Sl_m_↑). The gains in sex- bias are thus a result of subtle changes. SL_m_↓ lines, with increased size dimorphism and the smallest males, consistently express the highest number of sex-biased transcripts out of all lines (9,883 and 9,603, at FDR < 0.05, in replicates A and B, respectively; figures 2 and S2) . The SA treatment, despite a very consistent change in the phenotypic SSD (figure 1), shows the largest between replicate variation in the number of sex-biased transcripts (SA A and SA B, figures 2 and S2). The proportion of sex-biased transcripts that are shared by the SA replicate lines is nevertheless similar to other selection modes (e.g. replicate line overlap for male-biased transcripts: 36.9% in the SA, compared to 40.3%, 36.0%, 38.1% in the SL_m_↓, SL_f_↑ and SL_m_↑ modes respectively; female-biased: 39.5% in the SA, compared to 42.5% 41.2%, 41.6% in SL_m_↓, SL_f_↑ and SL_m_↑ modes respectively).

We thus conclude that the variation in the numbers of sex-biased transcripts is mostly caused by line-specific changes, and represent subtle gains and losses of sex-bias, with a higher number of male-biased transcripts unique to a specific line. We find no support for an transcriptome-wide relationships with sexual size dimorphism, but evidence that male-limited selection increases the number of sex-biased transcripts more readily and more consistently than SA or female-limited selection.

### How does the mode of selection affect the extent and parallelism of expression change, and cross-sex correlations in expression, in ancestrally sex-biased and unbiased transcripts?

PCA analysis of expression profiles shows that the first two principal components separate the sexes (1^st^ PC axis) and the selection lines (2^nd^ PC axis; figure S4) as expected based on their phenotypic differences. We next examine consistent patterns of differential expression (DE) across the replicate lines at all 19,373 transcripts, and compare the selection modes separately in each sex. We specifically test for gene expression consequences of body size evolution in the presence and absence of the genomic constraint under SA and sex- limited selection, respectively, on ancestrally sex-biased and unbiased transcripts (i.e. as defined in the C line). We first contrast the selection modes subjected to bi-directional body size selection on males: In the SL_m_↓ vs. SL_m_↑ contrast selection was for smaller (SL_m_↓) and larger (SL_m_↑) males through male-limited selection. Meanwhile in the SA vs. SL_m_↑ contrast, selection was also for smaller (SA) and larger (Sl_m_↑) males, but included SA selection. We thus examine the consequences of similar male phenotypic responses that were caused by different mode of selection (figure 1). We then contrast these modes to the control line (C) subjected to random selection on body size but with otherwise similar demographic changes.

### Expression divergence under SA selection is more constrained and sex-specific than under sex-limited selection

The DE transcripts show significant deviations in the distributions across chromosomes from those expected from all annotated transcripts, with fewer transcripts than expected on sex-chromosomes (table S1). There are overall fewer DE transcripts in the SA vs. SL_m_↑ compared to the SL_m_↓ vs. Sl_m_↑ contrast, and the sex-differences are also striking. Males have 1.6x more DE transcripts compared to females in the SL_m_↓ vs. SL_m_↑ contrast (2,842 and 1,820 DE transcripts in males and females, respectively, table S2), but the sex difference is especially large in the SA contrast where males show 2.4x more significant changes (1,550 and 650 DE transcripts in males and females, respectively, table S2). The fact that female expression changes are even more constrained by SA selection than male changes, leading to the greater sex-differences relative to male-limited selection, are well aligned with the lack of phenotypic response to SA selection in females (figure 1).

### Evidence for parallel changes at the level of transcripts but not functional GO categories

Splitting the transcripts by whether they are DE in the same direction in both contrasts or only in one of the contrasts, shows that, in males, male-limited selection (SL_m_↓ vs. SL_m_↑) has resulted in well over twice as many DE transcripts specific to this contrast (hereafter, *unique* SL_m_↓, N = 2,076) compared to changes specific to the contrast including SA selection (hereafter *unique* SA, N = 784). Many transcripts also overlap between the two, showing expression difference in the same direction (hereafter *parallel*, N = 759; table S3).

PCA analyses of line divergence based on the *unique* Slm↓, *unique SA*, and *parallel* transcripts (identified in the DE analysis) shows how the 2^nd^ PC axis separates the samples by selection line as expected, and explains ∼16-20% of the total variance (figure S5). Importantly, the C line, representing the ancestral phenotype (Kaufmann et al. 2021), clusters in between the selection treatment lines with smaller and larger individuals. This verifies how the expression differentiation between the body size selection lines is bi-directional in regards to the control line (figure S5). Specifically, the *unique* SL_m_↓ transcripts show divergent expression in the SL_m_↓ and SL_m_↑ away from the C along the PC2 in both sexes, while the SA cluster closer to the large SL_m_↑ (figure S5 A). The *unique* SA transcripts, in turn, show a clear bi-directional separation of the SA and SL_m_↑ lines, while both control and SL_m_↓ cluster in between (figure S5 C). Finally, PCA of *parallel* transcripts shows more limited divergence between the lines with small males (SL_m_↓ and especially SA) and the control, while SL_m_↑ is separated in its own cluster (figure S5 B), suggesting that most *parallel* transcript changes are caused by divergence due to large male selection (SL_m_↑).

We further identified the *unique* and *parallel* transcripts that show evidence of differential expression from the C line at the level of individual transcripts (hereafter referred to as *strict* lists). These include 267, 48, and 152 transcripts in the Sl_m_↓ contrast only (*strict unique* SL_m_↓), in both SL_m_↓ and SA (*strict parallel*), and in the SA only (*strict unique* SA), respectively. Thus, 10% (48/486) and 13% (48/361) show parallel expression change out of all the DE transcripts in males in the SL_m_↓ and SA contrasts to the C, respectively.

We tested for functional parallelism by comparing GO terms enriched among the *unique* SL_m_↓ and *unique* SA transcript sets (table S4 – unique L1, table S4 – unique SA, and table S4 – unique L1 – unique SA overlap) and found that the overlap of terms is not greater than expected by chance (from 100 bootstrapped overlaps with a random set of genes).

### Male-biased genes are disproportionately affected by body size selection

Several of our results suggest that body size evolution in our lines disproportionately affects the expression of male-biased transcripts. First, the transcripts DE in SL_m_↓ vs. SL_m_↑ contrast, for males, are enriched for male-biased transcripts compared to genome-wide expectations (*unique* SL_m_↓: X^2^ = 90.54, p < 0.001; *parallel* with SA: X^2^ = 54.35, p-value < 0.001; table 1A). These patterns remain when considering divergence from the control (*strict* sets of transcripts, table 1B), but are only statistically significant (at p < 0.05) in the *strict parallel* set (i.e. significantly DE also in the SA males). Second, the transcripts specifically upregulated in small males are more often male-biased than expected, regardless of the selection mode (i.e. both up-in-SL_m_↓ and up-in-SA, table S5), as are those upregulated in the large males (up-in-Sl_m_↑) in the SL_m_↓ v. SL_m_↑ contrast (table S5). Female-biased and unbiased transcripts are in turn under-represented among the male DE transcripts (table 1 and table S5).

**Table 1.** Associations with sex-bias among differentially expressed transcripts in males (see main text for description of the categories). Counts with asterisks indicate a significantly greater proportion than expected based on genome-wide proportions of sex-biased transcripts (shown in brackets in the column headers). Proportions out of row totals are given in brackets. **A)** Counts are the number of transcript in each category. Distributions are significantly different across rows (X^2^_6_ = 170.45, p < 0.001). **B)** Counts are the number of transcripts in each “strict” category (i.e. differences in expression also compared to the control (C) line. Distributions are significantly different across rows (X^2^_6_ = 28.19, p < 0.001)

|  | **Female-**<br>**biased**<br>**(0.23)** | **Male-**<br>**biased**<br>**(0.24)** | **Unbiased**<br>**(0.53)** | **Total** |
| --- | --- | --- | --- | --- |
| **A)** |  |  |  |  |
| *not-DE* | 3,649 (0.23) | 3,597 (0.23) | 8,508 (0.54) | 15,754 |
| *unique* SL<sub>m</sub>↓ | 438 (0.21) | 693* (0.33) | 945 (0.46) | 2,076 |
| *parallel* | 142 (0.19) | 273* (0.36) | 344 (0.45) | 759 |
| *unique* SA | 169 (0.22) | 179 (0.23) | 436 (0.56) | 784 |
| **B)** |  |  |  |  |
| *not DE* | 652 (0.21) | 1,011 (0.32) | 1,489 (0.65) | 3,152 |
| *strict unique*<br>SL<sub>m</sub>↓ | 55 (0.21) | 72 (0.27) | 140 (0.52) | 267 |
| *strict parallel* | 4 (0.08) | 27* (0.56) | 17 (0.35) | 48 |
| *strict unique* SA | 38 (0.25) | 35 (0.23) | 79 (0.52) | 152 |

The expression patterns of sex-biased transcripts in females of these lines are more variable, reflecting their different patterns of phenotypic divergence. Overall, distributions of sex-biased transcripts are different across the *unique* SL_m_↓*, unique* SA, and *parallel* transcript categories also for females (X^2^_6_ = 144.56, p-value < 0.001, table 2). Compared to the expected distribution, transcripts changing in the females of the SL_m_↓ vs. SL_m_↑ contrast in response to selection for small males (*unique* SL_m_↓) are enriched for male-biased but also female-biased transcripts (X^2^_2_ = 124.25, p < 0.001; table 2), and those specifically upregulated in the smaller SL_m_↓ females are disproportionately female-biased (up-in-SL_m_↓, table S5). Transcripts changing in parallel with SA selection in females have similarly more male-biased but fewer female-biased transcripts than expected (*parallel*: X^2^_2_ = 7.45, p = 0.02). Among those transcripts DE only in the SA females, there is no evidence for an excess of any sex-biased category (*unique* SA: X^2^_2_ = 3.81, p = 0.15; table 2). Even the transcripts upregulated in females of SA lines, where selection acted towards larger females (up-in-SA) are, if anything, lacking in female-biased transcripts (table S5). Those transcripts with higher expression in females due to selection for larger males (up-in-SL_m_↑) are disproportionately male-biased and/or unbiased, depending on the contrast (table S5). Thus, while male-biased expression is readily altered by all modes of selection and in both sexes, female-biased transcripts are disproportionately affected only in females and only due to male-limited selection.

**Table 2.** Associations with sex-bias among differentially expressed transcripts in females (see main text for details). Counts with asterisks indicate a significantly greater proportion than expected based on genome-wide proportions of sex-biased transcripts (shown in brackets in the column headers). Proportions out of row totals are given in brackets. Distributions are significantly different across rows (X^2^_6_ = 144.56, p < 0.001).

|  | **Female-biased**<br>**(0.23)** | **Male-**<br>**biased**<br>**(0.24)** | **Unbiased**<br>**(0.53)** | **total** |
| --- | --- | --- | --- | --- |
| *not-DE* | 2,674<br>(0.17) | 2,610<br>(0.16) | 10,728<br>(0.67) | 16,012 |
| *unique SL<sub>m</sub>↓* | 496*<br>(0.32) | 458*<br>(0.29) | 615<br>(0.39) | 1,569 |
| *parallel* | 42<br>(0.17) | 76*<br>(0.31) | 131<br>(0.53) | 249 |
| *unique SA* | 94<br>(0.19) | 120<br>(0.24) | 276<br>(0.56) | 490 |

### Cross-sex correlation in expression changes is modified by sex-bias but not by selection mode

Although different numbers of transcripts are DE in the sexes, especially in the SA lines (see above), the expression changes across the transcripts (significant in either sex) are collectively highly correlated between males and females in both SL_m_↓ and SA contrasts (to both SL_m_↑ and C; figure 3, figure S7). As we predicted (figure 1), the cross-sex correlation coefficients are significantly lower for sex-biased than for unbiased transcripts (i.e. non-overlapping confidence intervals) for most transcript categories (figure 3, figure S7). We find that the mode of selection has only a marginal and inconsistent effect on the correlations, and no support for the prediction that SA selection would have resulted in a lower net correlation between the sexes, despite the phenotypic evolution occurring only in males (figure 3). Taken together, the DE and correlation analyses suggest that SA selection has caused more parallel (with SL_m_↓) and a subtly greater number of changes in the males compared to females, rather than major differences and in different sets of genes.

**Figure 3.**
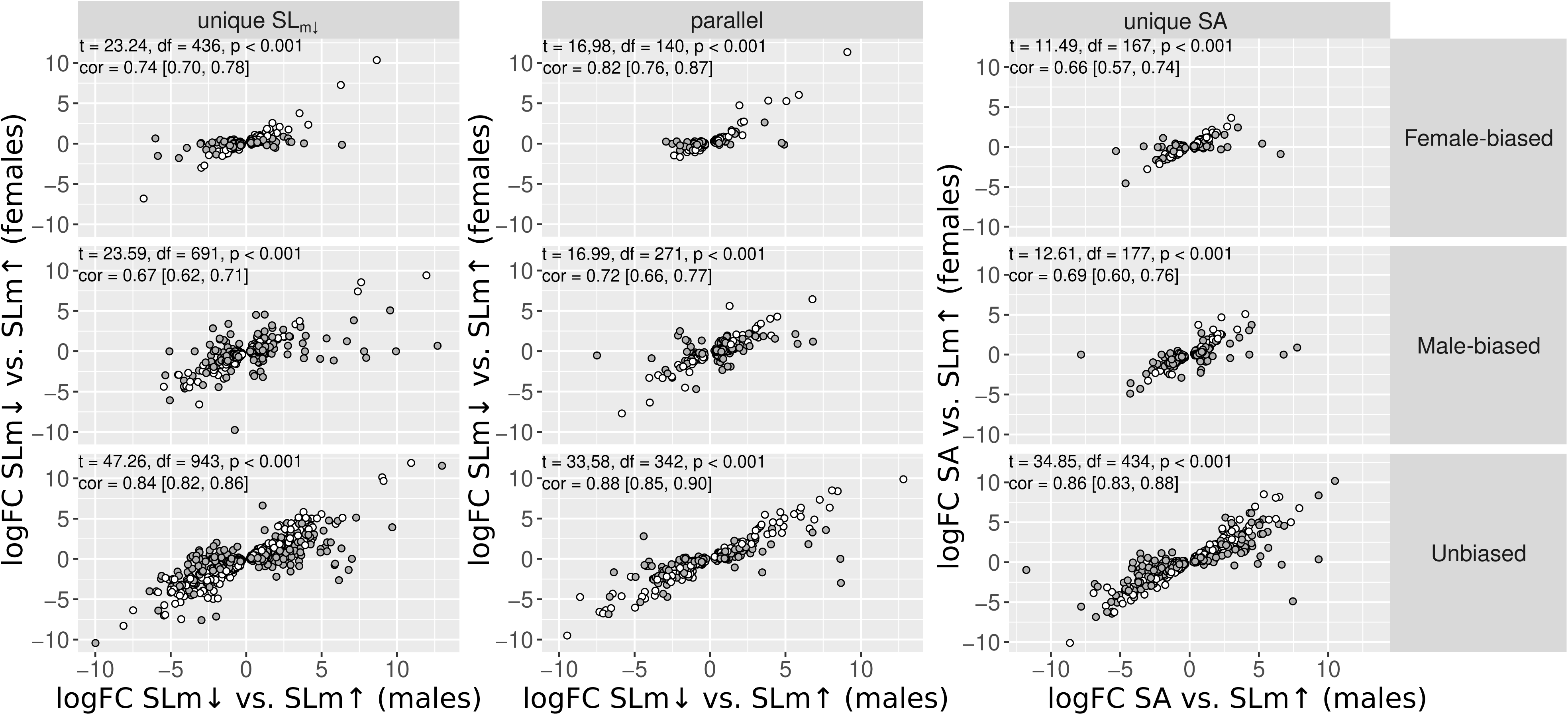
logFC expression between SL_m_↓ and SL_m_↑, or SA and SL_m_↑, for males and females. Data are split by whether males show a unique response in the SL_m_↓ vs. SL_m_↑ contrast, a unique response in the SA vs. SL_m_↑ contrast, or a parallel response in both, and by sex-bias status. Points are coloured by whether both of the sexes show the same significant expression difference (white) or only one of them (grey). Inset text gives results of Pearson’s correlation tests as well as correlation coefficients for each panel. Note that the axis labels are different for the final column of panels. The logFC values for males and females of the parallel transcript class in the SA vs. SL_m_↑ contrasts is given in the supplementary materials (figure S6).

### The sex under selection influences the extent of expression changes but not the patterns of sex-bias or the cross-sex correlations

Our selection lines allow us to also test how a similar selection mode, but acting on a different sex, affects expression changes, by comparing lines with female- (SL_f_↑) and male- limited (SL_m_↑) selection for larger size to the selection line for smaller (male) size (SL_m_↓). In the SL_m_↓ vs. SL_f_↑ contrast, females had opposite phenotypic responses, but due to selection on different sexes. In the contrast SL_m_↓ vs. SL_m_↑, females also had opposite phenotypic responses, but in this case entirely due to selection on males. We further compare female expression in these lines with the C line (producing the *strict unique* SL_f_↑, *strict unique* SL_m_↑, and *strict parallel* sets for females in these contrasts).

Selection solely limited to males (i.e. SL_m_↑ vs. SL_m_↓ contrast) has affected expression considerably more than when selection also targeted females (SL_m_↓ vs. SL_f_↑), with the greatest difference seen in male expression changes (table S6, table S7). Notably, unlike under purely male-limited selection where significantly more transcripts were affected in males (table S2), a similar number of transcripts have changed in each sex when selection was limited to females (686 and 650 DE transcripts in females and males, respectively, table S6) and where the phenotypic evolution in the sexes was also largely similar (figure 1). This is in stark contrast with the male-limited and SA selection modes with sex-specific phenotypic changes. The expression changes between the sexes are also highly correlated, as in the other selection modes. Coefficients for the sex-biased and especially female-biased transcripts are significantly lower than for the unbiased transcripts (figure S8), as also expected.

There are 366 and 288 transcripts that respond in parallel in both contrasts which are 10% and 12% of all the DE transcripts in these contrasts in females and males, respectively (table S7). The overlap of parallel transcripts in females is thus very similar to under selection targeting only males or antagonistically both sexes (11%, table S3). These overlapping transcripts constitute 39% and 56% of the changes occurring in females and males, respectively (table S8), but 11% and 13% of the corresponding changes under solely male- limited selection, reflecting the overall greater divergence under the latter. Only very few DE transcripts differ also from the control due to female-limited selection for larger size (n = 46; *strict unique* SL_f_↑), while 327 transcripts have changed only due to male-limited selection for larger size (*strict unique* SL_m_↑). Notably, 19 transcripts are DE in both contrasts (*strict parallel*, 29% and 5% out of all significant *strict* transcripts in the contrasts of SL_f_↑ and SL_m_↑ to the control, respectively).

Female-biased transcripts are generally under-represented, with the exception (again) of those upregulated in the small females of SL_m_↓ lines in both contrasts (table S8). DE transcripts upregulated due to female-limited selection (up-in-SL_f_↑) are more commonly male-biased in both males and females (table S8). The transcripts *unique* to male-limited selection for large size, and those in *parallel* with the female-limited selection, show an excess of male-biased transcripts (table S9 *c.f.* table 1), while there are no deviations from expectations for *unique* changes in the female-limited selection (SL_m_↓ vs. SL_f_↑; table S9). However, the upregulated DE transcripts of the SL_f_↑ are more commonly male-biased in both males and females (up-in-SL_f_↑; table S8), while female-biased transcripts are under- represented (with the exception, again, of those upregulated in the small females of SL_m_↓ lines in both contrasts; table S8). Out of the few SL_f_↑ transcripts that differ from the control, only three are female-biased in expression, while 14 are male-biased, a significant deviation from expectations (X^2^ = 6.88, p-value = 0.03).

## Discussion

We experimentally tested how adult gene expression changes in response to different modes of sex-specific selection on body size. We took advantage of the known selection history and phenotypic evolution in our artificial selection lines, and compared the effects of sex-limited (SL) and sexually antagonistic (SA) selection that has acted in the same direction on males, but not females, as well as SL selection that has operated similarly but on opposite sexes. We also utilized a single replicate of a control line subjected to similar demographic changes as the selection lines (see Materials and Methods). We find that thousands of transcripts are differentially expressed between the selection modes in both sexes with the vast majority of changes at autosomal loci. PCA analyses indicate that the selection lines show expected expression divergence from the control line. Male- and female-biased transcripts are differently impacted by SA and SL selection, and collectively show less correlated responses between the sexes, than unbiased transcripts, as we predicted. The degree of sex differences in significant expression changes are somewhat aligned with sex-differences in phenotypic responses (figure 1), being greatest in the SA selection lines and smallest in the female-limited selection lines. We do not, however, find support for a transcriptome-wide association between sex-bias and sexual dimorphism. Below we discuss in detail the extent to which expression changes reflect phenotypic evolution in the sexes and are modulated by the mode of selection, in the light of our initial study questions.

### Sexual size dimorphism, sex-biased gene expression, and sexual conflict

Two decades of genome-wide studies of gene expression across taxa have revealed extensive gene regulatory differences between the sexes. The conceptual framework of sexual conflict assumes that such regulatory decoupling evolves under SA selection to alleviate sexual conflict. The direction of sex-bias is therefore thought to reflect the different fitness optima in each sex in gene expression (Ellegren and Parsch 2007; Ingleby et al. 2015; Pennell et al. 2024), and in phenotypes, predicting a positive covariation between the two (Pointer et al. 2013; Harrison et al. 2015; Mank 2017; Wright et al. 2018; Cheng & Houle 2020). To date, the empirical evidence remains inconclusive (Pointer et al. 2013, Harrison et al. 2015; Toubiana et al. 2021 *vs*. e.g. Khila et al. 2012; Scharmann et al. 2021). Here, we tackled this question by studying how the evolution of sexual dimorphism in a single trait affects gene expression, as a population-level process. We followed a common approach in the field and used bulk RNA samples to study expression as a molecular phenotype, and to make our study comparable (but see e.g. Price et al. 2022).

We found no support for the prediction of a transcriptome-wide association between sex-biased expression and evolved sexual size dimorphism (figure 2 and figure S3). The majority of analysed loci are similarly unbiased or sex-biased across the lines, but there is also variation in their number and proportion across the lines demonstrating how sex-bias can be lost and gained rapidly in different populations on small evolutionary timescales (figure 2 and S3). This turnover was largely specific to each replicate line rather than selection mode, and may therefore reflect more drift than selection.

While our data does not support the prediction that unbiased genes consistently obtain sex-bias when selection increases phenotypic sexual dimorphism, we do find that both SA and sex-limited selection disproportionately alter expression at sex-biased loci. Male-biased transcripts are enriched among expression changes and more commonly upregulated in the males of all the lines evolving towards the male-optimal smaller size (SA and Sl_m_↓; and even in females but in a small subset of loci). SA and SL_m_↓ lines not only evolved significantly smaller males, but also increased sexual size dimorphism due to no (under SA) or slower (under SL_m_↓) response in females (Kaufmann et al. 2021; see figure 1). However, male-biased transcripts are upregulated even in the males due to male-limited selection towards larger size, which decreased sexual dimorphism, as well as in the female-selection lines, although the evidence here is much weaker. Together with the more frequent gains of novel male-bias in the selection lines, the differential expression analyses thus reveal that expression of male-biased transcripts evolves more rapidly, and consistently, than of female-biased transcripts, which signifies selection response rather than drift. This micro- evolutionary evidence supports the frequent finding in comparative studies of fast expression evolution of male-biased genes in much larger timescales (Parisi et al. 2003; Meiklejohn et al. 2003; Martin et al. 2013; Harrison et al. 2015; Yang et al. 2016). Our lines also show that life-history evolution by natural selection contributes to this phenomenon that has mostly been attributed to sexual selection or drift, echoing our recent findings in another seed beetle species (Immonen et al. 2023).

According to the chief assumption of positive relationship between molecular and phenotypic sexual dimorphism, we should have observed mirrored changes in female-biased transcripts, towards increased female-biased expression in lines that have been selected towards female-optimal larger size (i.e. SA and SL_f_↑). Instead, we observed that female-biased transcripts are most numerous, and most commonly differentially expressed in the females of the small male selection lines (SL_m_↓), which overall possess the highest number of sex-biased transcripts. In contrast, female-biased transcripts are not particularly affected by either SA or female-limited selection in our study (in either sex). The strength of selection tends to increase with expression level and it is therefore generally assumed that sex-specific selection will be stronger on the highly expressed genes of the targeted sex (Mank 2017; Tosto et al. 2023). The deficit of changes in female-biased transcripts contradicts this and shows that evolution towards female-optimal phenotype does not automatically feminise the transcriptome as is commonly expected in other studies (Hollis et al., 2014; Immonen et al. 2014, Veltsos et al. 2017; Parker et al., 2019).

Our data thus demonstrate that sex-limited selection is capable of modifying sex-biased expression to an even greater extent than SA selection, despite the latter increasing SSD more than SL selection for similarly small males (Kaufmann et al. 2021; figure 1; figure 2). This is not surprising given that in theoretical work it has long been recognised that sexual dimorphism can evolve even under sexually concordant selection when genetic variances are asymmetric in the sexes (Fisher 1930; Lande 1980; Bonduriansky & Chenoweth 2009; Gosden et al. 2012; Wyman et al. 2013; Cheng & Houle 2020; Houle & Cheng 2021). To understand these results, it is useful to consider the main components that affect expression evolvability. First, the direct expression response to body size selection in any given gene is determined by its transcriptional role in regulating body size, and the amount of genetic variation underlying its expression in the sex under selection (e.g. Cheng & Houle 2020, Houle & Cheng 2021). Based on our results we can speculate that, as a group, the autosomal male-biased loci studied here may harbour more genetic variation in the elements underpinning regulatory variation (e.g. transcription factors and their binding sites, chromatin remodelling, epistatic trans effects from sex chromosomes) thus facilitating their relatively higher expression evolvability. Higher genetic variation for expression of male-biased genes has been detected in other systems too (Allen et al. 2018).

Second, because genes are often expressed as part of gene regulatory networks, they can also evolve indirectly via cross-gene genetic correlations (Houle & Cheng 2021), but may also be prevented from evolving by these correlations. More pleiotropic regulatory networks may constrain direct response of individual genes. Female-biased genes show a wider expression breadth across tissues in *C. maculatus* (Immonen et al. 2017), as in many other systems, which suggests that female-biased genes may be more commonly constrained by pleiotropy (Allen et al. 2018; Dean & Mank 2016). Their lack of response when selection targeted females (i.e. in SA and SL_f_↑), combined with increased expression only when selection was relaxed on females (SL_m_↓), supports this view.

Third, indirect responses also occur because of the positive cross-sex genetic correlations, estimated with the *r_m,f_,* which is the primary cause of expression changes in the sex *not* under selection in our SL selection lines. Expression *r_m,f_* is negatively associated with evolvability of sex differences (Houle & Cheng 2021). A high and positive *r_m,f_* should thus limit direct expression responses to SA selection, but not to SL selection, where it can generate a greater net response across both sexes due to direct and indirect changes. This is also what we see: the expression changes under SA selection are overall far fewer than under SL_m_↓ selection. This is especially so in females: SA line males show over 2.4 times more significant changes than females, while SL_m_↓ line males only 1.6 times more (table S2). The large sex difference reflects the phenotypic changes well (Kaufmann et al. 2021) and suggests that SA selection has affected the sex-specific regulatory architecture underlying body-size such that SA males were able to change relatively more than females. Under male-limited selection, male size evolved even slightly further than under SA, and female size showed a strong correlated response (Kaufmann et al. 2021), thereby contributing to greater erosion of genetic variance in body size under this selection mode compared to the SA (Kaufmann et al. 2023a). The greater number of changes observed in the SL_m_↓ lines thus reflects this overall greater divergence.

Despite the large sex difference in the numbers of significant transcripts in the SA lines, the expression changes are highly correlated between the sexes, which suggest that the sex differences are subtle. The changes in the SA females are predominantly in unbiased transcripts (table 2), which show a generally higher cross-sex correlation in expression changes than sex-biased transcripts, as we predicted. There is no difference however in the strength of this correlation between the selection modes (figure 2). Similarly, although we find good evidence that the cross-sex correlations are lower for sex-biased transcripts, and male-biased transcripts are disproportionately affected by SA selection in males, the correlations are no different between SA and SL selection treatments. Indeed, neither selection mode nor the sex under selection has any effect on the strength of the correlations (figure S8). The general pattern of lower correlated responses in sex-biased transcripts in our lines indicates that the underlying *r_m,f_*in expression is relatively lower in sex-biased loci, in accordance with studies in *Drosophila*, where the association between the *r_m,f_* and sex-bias is overall negative (Griffin et al. 2013; Allen et al. 2018; Houle & Cheng 2021). Taken together, the correlated evolution in the sexes we observe suggests that while small sex differences can clearly evolve quite readily, the sexes are nevertheless constrained by their shared expression architecture, which indicates that the potential for sexual conflict is rampant.

### Parallel changes and consistent targets of selection

The extent to which parallel gene expression changes underlie parallel phenotypic changes has been tested both in wild populations (Jeukens et al. 2010; Zhao et al. 2015; Yeaman et al. 2016) and in replicate populations in experimental evolution studies (Cooper et al. 2003; Immonen et al. 2014; Veltsos et al. 2017; Hsu et al. 2020; Veltsos et al. 2022; Thorhölladottir et al. 2023; Mishra et al. 2024; Ghalambor et al. 2015; José Rivas et al. 2018; McGirr & Martin 2018; Fischer et al. 2021; Jacobs et al. 2020, Immonen et al. 2023). These studies have found both repeatable and idiosyncratic changes. In typical experimental evolution studies, parallelism is inferred from consistent responses across similarly treated replicates, which is the foundation for detecting changes due to selection, as opposed to drift. Our differential expression analyses also rely on this consistency among the selection line replicates, but expands the question to whether it is possible to detect parallel changes when phenotypic evolution is similar but the mode of selection, or the sex under selection, is not. Our results indicate that the answer is “yes” to both. The SA and SL selection for small males affect the expression of the same transcripts in about a 10^th^ of the cases (i.e. parallel changes in 13% and 10% of all the significant loci in SA and SL_m_↓ male contrasts to both control and SL_m_↑, respectively, i.e. *strict parallel*). When comparing similar selection modes but acting on a different sex, parallelism is increased to 29% (i.e. out of all changes from the control line due to sex-limited selection on females for larger size), although fewer transcripts are different from the control line. What causes differences in evolvability under similar selection but acting on the different sexes is the fact that the genetic architecture (G matrices) for expression are never expected to be identical in the sexes (Houle & Cheng 2021). Moreover, lower parallelism when comparing the different selection modes can be expected due to the fact that the genomic constraint imposed by SA selection is restrictive in the changes that can occur. These should primarily involve loci with a net positive effect on fitness across both sexes, which is not required under male-limited selection.

Adult body size is a complex polygenic trait, and polygenic adaptation is expected to show low genetic parallelism and greater genetic redundancy (Barghi et al. 2020; Schlötterer 2023). The degree of parallelism can increase at the higher levels of biological organisation, from gene regulation to molecular pathways and physiological processes (Edelman et al. 2001). Here, we find no evidence that the overlap in enriched GO terms between *unique* SA and *unique* SL_m_*↓* transcript sets is greater than expected by chance, suggesting that distinct processes are impacted when selection has changed male, but not female size. The selection that changes adult size ultimately targets genes that regulate growth in the embryonic and/or juvenile stages. In our lines, the body size evolution seems to be produced mainly by differences in initial embryo size and/or in growth rate during juvenile stages (*unpublished data*). The gene regulatory differences in adults thus reflects developmental inertia at causal loci or downstream pleiotropic effects of these changes. A key growth pathway across taxa – insulin/insulin-like signalling and target of rapamycin (IIS/TOR) – is responsible for growth by regulating developmental proliferation of cell size and number (Nijhout et al. 2014; Texada et al. 2020), but also for maintaining the size/number of cells in adult organs by modulation of cell growth, division and death (Yang & Xu 2011). In accordance, we find that *“Toll signalling pathway”* GO terms are enriched among transcripts showing *parallel* expression differences in SA vs. SL_m_↑ and SL_m_↓ vs. SL_m_↑ contrasts (table S4 - parallel). The terms associated specifically with changes due to SA selection (i.e. *unique* differences from control and large-male selection, table S4) include many that are involved in growth rate regulation (unbiased transcripts), metabolic processes (male-biased transcripts) and mitosis (female-biased transcripts), while those associated with small size due to male-limited selection (i.e. *unique SL_m_↓* differences, table S4) include various lipid biosynthetic processes (unbiased transcripts), regulation of metabolic processes (male- and female- biased transcripts), and immune system processes (female-biased transcripts). However, a major caveat here is that only ∼37% of all transcripts have GO terms (see Materials and Methods), and this percentage is highly variable across the sets of DE genes (10-61%).

## Conclusions

Our results demonstrate that the sexually dimorphic transcriptome evolves rapidly under sex-specific selection. We show that the expression changes are less correlated across the sexes at sex-biased than at unbiased loci, regardless of the mode of selection. The strength of this correlation is surprisingly robust to the mode of selection and whether the loci are male- or female-biased. We find no evidence for transcriptome-wide correlation between sex-biased expression and sexual size-dimorphism across selection lines. However, male-biased autosomal loci are enriched among transcripts showing significant expression changes in the most sexually dimorphic lines. The lack of clear association with sexual dimorphism is an important result because it is frequently an implicit expectation in comparative studies of sex-biased genes, which have found very mixed support for it (Pointer et al. 2013; Harrison et al. 2015; Toubiana et al. 2021; Scharmann et al. 2021; Price et al. 2022). Our results also indicate that inferring the selection history on each sex from patterns of sex-biased gene expression is highly questionable. SA selection affects sex-biased expression, but sex-limited directional selection can do so too, and even to a greater extent (female-biased transcripts in this case). Our results suggest that expression at male- and female-biased transcripts is not affected in the same way or to the same degree, indicating a strong dependence on the specific loci underlying the traits under selection, and the genetic architecture of gene expression at these loci. This is consistent with SA selection constraining gene expression changes relative to sex-limited directional selection. This further suggests that rapid expression evolution of sex-biased genes reported between populations and species in some studies may more likely imply an absence of intra-locus sexual conflict rather than the presence of it.

## Materials and Methods

### Study organism, selection lines, and sequencing

*Callosobruchus maculatus* seed beetles are a subtropical and tropical pest of bean seeds with a cosmopolitan distribution. It has a rapid life cycle of approximately three weeks. Females lay eggs on the surface of beans that serve as food resource and environment for the larval development which, in our standard laboratory conditions, lasts approximately 25 days (at 29C°, 50% relative humidity, and a 12h/12h light/dark cycle). The beetles are aphagous capital breeders and adults complete their reproductive cycle relying solely on the larval resources. The adult body mass acquisition is thus exclusively a larval developmental process. *C. maculatus* has 18 autosomes and XY sex chromosomes.

In this study, we assessed experimentally how different modes of sex-specific selection – sexually antagonistic (SA) or sex-limited (SL) – affect gene expression in adult beetles. The selection lines utilised here have been described in detail in Kaufmann et al. (2021). All lines originate from a single wild collected population (from Lome, Togo). The selection lines were created by applying 10 generations of artificial family-level selection either only on males, towards smaller or larger size (SL_m_↓, SL_m_↑ respectively), only on females, towards larger size (SL_f_↑), or sexually antagonistically (SA) on both sexes for larger females and smaller males (i.e. increasing the naturally occurring sexual size dimorphism). Each mode of selection was independently replicated twice. We also use here a single replicate of the control line (C) subjected to similar demographic changes as the selection lines but to relaxed (random) selection on size. The C line has retained the ancestral patterns of larval growth and adult body size (line Cb in Kaufmann et al. 2021, 2023a). Our quantitative genetic analysis (Kaufmann et al. 2023a) confirms that this control line has largely maintained the ancestral genetic variance for body size, this is also confirmed by our Pool-seq analysis (Zwoinska et al. *in prep.*). Additionally, we rely mostly on contrasts of the fully replicated experimental treatment lines, using C lines as a follow-up analysis to determine whether divergence is likely to be bi- or uni-directional from C. This strategy assures that we capture divergence between the extremes of body size evolution with minimal effect of drift. Thus, we are confident that a single C line replicate is representative of the control lines, although it remains possible that subtle idiosyncratic drift effects may have occurred, which will be missed.

Due to logistic limitations, it was not possible to start the gene expression experiments directly after selection was terminated. We therefore established isofemale lines from each selection line to limit the impact of relaxed selection. Eight isofemale lines per selection line (each started by a single full-sib family) were kept under standard rearing conditions for approximately a year (∼15 generations). To create the samples for RNA-seq, the isofemale lines were first density controlled for two generations (by limiting oviposition window to 24h, on a surplus of beans). Only beans with a single egg (and thus no larval competition) were used to collect virgin adults for the experiment. We then chose six isofemale lines and made three inter-line crosses. From the F1 progeny of these crosses, we picked five males and five females per cross to produce pooled samples by sex. The adult beetles were snap-frozen in liquid nitrogen 24h after eclosure. Each sequenced sample thus represents an average profile of five individuals from two isofemale lines. For each selection line, we produced three such pools per sex, resulting in six samples representing each selection line. The total number of samples sequenced was 54. Total RNA was extracted from whole bodies, using Qiagen RNeasy, according to manufacturer’s instructions. The RNA was checked for quality and quantity using NanoDrop, Qubit and BioAnalyzer. TruSeq stranded mRNA libraries were prepared for each sample including poly-A selection to enrich for mRNA. Paired-end sequencing for read lengths of 150bp was performed on a single NovaSeq 6000 S4 flowcell. Library preparation and sequencing was performed by the SNP & SEQ Technology Platform of SciLifeLab in Uppsala.

### RNA-seq read pre-processing, and expression quantification

We used the most recent genome assembly (made from the same population as the selection lines) and the associated annotation (Kaufmann et al. 2023b). RNA-seq reads were first trimmed to identify and remove TruSeq adapters with cutadapt (v. 4.0; Martin 2011). After trimming, any reads <75bp in length were discarded. We quality checked against any putative remaining rRNA reads in the libraries, using SortMeRNA (v. 4.3.3; Kopylova et al. 2012) to identify reads that mapped to a set of Coleopteran rRNA sequences, obtained from NCBI (n = 2,646 sequences; https://www.ncbi.nlm.nih.gov/) and SILVA (n = 13,768 sequences; Pruesse et al. 2007; Quast et al. 2013). These sequences were clustered with cd-hit (v. 4.8.1; Li et al. 2006; Fu et al. 2012) by similarity of 95%. We retained all reads that did not map to this curated database of rRNA sequences. We used Salmon (v. 1.6.0; Patro et al. 2017; Srivastava et al. 2020) in selective alignment mode to quantify the expression of all transcripts in the *C. maculatus* transcriptome. We engaged the salmon options --gcBias -- seqBias, and --posBias.

For a subset of samples (from the control line), we mapped reads also to the genome assembly using STAR (v. 2.7.9a; Dobin et al. 2013), and used ExplorATE (Femenias et al. 2022; available from: https://github.com/FemeniasM/ExplorATE_shell_script; commit: ecebe32) to quantify the contribution of TE activity to the mRNA samples (see the Supplementary Materials: Supplementary Methods).

Prior to differential expression analysis we excluded transcripts on contigs shorter than 100kb (n = 5,977; 15%) to limit our analysis to the most reliably assembled and annotated portion of the genome (Kaufmann et al. 2023b). We also removed transcripts on the Y chromosome (n = 437) to avoid confounding results due to multi-mapping to autosomal homologs. We additionally performed minimal filtering of very lowly expressed transcripts by removing any transcripts with overall mean counts < 3 across all samples (n = 13,513). Following this filtering, we were left with expression estimates from 19,373 transcripts in total. We also conducted a principle component analysis (PCA) with the prcomp() function in R (v. 4.3.3; R Core Team 2024) to establish the overall patterns of gene expression across sexes and selection lines. For the PCA we additionally removed transcripts with no variance in the Transcripts Per Million (TPM) estimates across samples (n = 494), as these are uninformative in PCA.

### Is there an association between sexual size dimorphism and transcriptome-wide sex-bias?

#### Sex-biased transcript expression

We first tested for differences in expression between the sexes within each selection line and replicate. To account for the structure in the data arising from male and female pools being linked by the cross that produced them, we included a cross term as a fixed effect in the model:

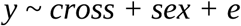

All the p-values were adjusted for multiple testing using the Benjamini-Hochberg correction and control FDR at p_adjusted_ < 0.05. For each selection treatment, we computed the number of sex-biased transcripts, as well as their relative proportions, across chromosome categories (autosomes, X-chromosome,). We then tested for an association between the number and proportion of sex-biased transcripts on each chromosome type and the degree of sexual size dimorphism in each selection treatment (the mean SSD across the two replicate selection lines, data from Kaufmann et al. 2021). We also tested for an association between the magnitude of sex-bias (median log(fold-change) between males and females across all sex-biased transcripts) and the degree of sexual size dimorphism in each selection treatment. We consider female-biased (log(fold-change) > 0) and male-based (log(fold-change) < 0) transcripts separately.

We also formally compared the degree of sex-bias between the C line and each selection treatment line replicate in a joint model by testing a sex-by-line interaction term. For this analysis we created a sex:line composite factor for each sample and fit the model:

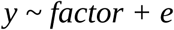

In these analyses we did not have enough data to include a cross effect. Instead, we defined a specific contrast set always within sex:

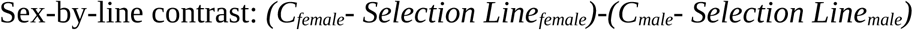

Thus, if a test of this contrast shows evidence for an effect, then the difference between C and the selection line differs by sex.

### How does the mode of selection affect the extent and parallelism of expression change, sex-biased gene expression, and cross-sex correlations in gene expression?

#### Gene expression differences across selection treatments

We next sought to identify transcripts showing expression differences associated with the selection treatments. In our analytical pipeline, we were interested in finding the changes associated with divergence in male size under *1)* male-limited selection (SL_m_↑ vs. SL_m_↓) and *2)* SA selection (SL_m_↑ vs. SA), and how these two contrasts may compare given the similarities in male but differences in female phenotypic evolution. *3)* We also tested whether the sex under selection matters by contrasting also the selection lines for large size via females (SL_f_↑ vs. SL_m_↓). To evaluate the degree of sex-specific responses to selection we conducted these tests separately for each sex, and tested how correlated the expression differences were between the sexes, using spearman rank correlation tests.

Because replicate lines were assigned to selection treatments independently, they should be treated as unique. To identify changes across the replicates in a given treatment, we constructed a contrast to test for an average difference between selection treatments, such that:

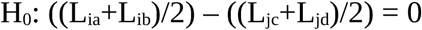

Where *i* and *j* are different selection treatments, and *a, b, c,* and *d* are independent replicate lines. Additionally, for the transcripts of interest (see Results) we tested for differences between the selection treatment lines and the control line, C:

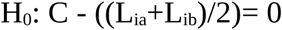

All differential expression analysis were performed with the edgeR package (v. 4.4.2; Robinson et al. 2010; Chen et al. 2025) in R (v. 4.4.2; R Core Team 2024) with the default library normalization and transcript filtering procedures. As a reference in comparative analyses between the selection lines (see below), we used sex-biased transcripts called in the control line (C). Although the extent to which the sex-bias state in the C line reflects the ancestral state is not known (due to no data from the ancestral population), several lines of evidence suggest they are in close accordance. The control line has retained most ancestral genetic variance in body size (Kaufmann et al. 2023a), shows limited genome-wide divergence in allele frequencies from the ancestral population (Zwoinska et al. *in preparation*), and shows no body size divergence from the ancestral population in either sex (Kaufmann et al. 2021).

For all tests, we controlled the FDR at p_adjusted_ < 0.05.

We used the R package ViSEAGO (v. 1.15.0; Brionne et al. 2019) for GO term enrichment analysis, setting all transcripts as the background set and using the “Biological Process” ontology terms. We note that ∼37% of annotated genes received GO terms, these enrichment analyses must be interpreted with that in mind. We tested for an overlap in enriched GO terms using a bootstrapping procedure. We performed 100 boostrap samples of *n* random transcripts, where *n* was always the equivalent number of transcripts to the set being compared.

## Acknowledgements

Sequencing was performed by the SNP&SEQ Technology Platform in Uppsala. The facility is part of the National Genomics Infrastructure (NGI) Sweden and Science for Life Laboratory. The computations were enabled by the Swedish National Infrastructure for Computing at UPPMAX (by projects SNIC 2022/5-83; 2022/6-37; 2023/5-111; 2023/6-65) The work was also supported by grants from the Swedish Research Council (*Vetenskapsrådet*) [no. 2019-05038 and 2023-04869], and Carl Tryggers foundation to EI [grants no. CTS-18:163 and CTS-19:155], by grants from the Swedish Research Council (*Vetenskapsrådet*) [grant no. 2022-03701], and Marie Skłodowska-Curie Actions (European Commission) to RAWW [grant no. 101061664], by grants from The Nilsson-Ehe Endowment and Stiftelsen Lars Hiertas Minne [grant nos. 40933 and FO2019-0329] to JMH, and by a grant from the Birgitta Sintring Foundation [grant no. S2024-0007] to MKZ. We also thank the three anonymous reviewers and the associate editor for comments that have greatly improved the manuscript.

## Data availability

Data and analysis scripts have been deposited in a zenodo archive with doi: 10.5281/zenodo.13919165. Raw reads have been deposited at NCBI with BioProject accession: PRJNA1178449

## References

1. Arnqvist, G. & Tuda, M. (2010). Sexual conflict and the gender load: correlated evolution between population fitness and sexual dimorphism in seed beetles. Proceedings of the Royald Society B: Biological Sciences. 277: 1345–1352.

2. Allen, S.L., Bonduriansky, R. & Chenoweth, S.F. (2018). Genetic constraints on microevolutionary divergence of sex-biased gene expression. Philosophical Transactions of the Royal Society B: Biological Sciences. 373: 20170427.

3. Austad, S.N. & Fischer, K.E. (2016). Sex differences in lifespan. Cell Metabolism. 23: 1022–1033.

4. Barghi, N., Hermisson, J. & Schlötterer, C. (2020). Polygenic adaptation: a unifying framework to understand positive selection. Nature Reviews Genetics. 21: 769–781.

5. Berger, D., Berg, E. C., Widegren, W., Arnqvist, G. & Maklakov, A. A. (2014). Multivariate intralocus sexual conflict in seed beetles. Evolution 68: 3457–3469.

6. Berger, D., Martinossi-Allibert, I., Grieshop, K., Lind, M.I., Maklakov, A.A. & Arnqvist, G. (2016). Intralocus sexual conflict and the tragedy of the commons in seed beetles. The American Naturalist. 188: 98–112.

7. Bonduriansky R. & Chenoweth S.F. (2009). Intralocus sexual conflict. Trends in Ecology and Evolution. 24: 280–288.

8. Brionne, A. Juanchich A. & Hennequet-Antier C. (2019). ViSEAGO: a Bioconductor package for clustering biological functions using Gene Ontology and semantic similarity. BioData Mining. 12:16.

9. Chen, Y., Chen, L., Lun, A.T.L., Baldoni, P.L. & Smyth, G.K. (2025). edgeR v4: powerful differential analysis of sequencing data with expanded functionality and improved support for small counts and larger datasets. Nucleic Acids Research. 53: gkaf018.

10. Cheng, C. & Kirkpatrick, M. (2016). Sex-specific selection and sex-biased gene expression in humans. PLoS Genetics. 12: e1006170.

11. Cheng C. & Houle D. (2020). Predicting multivariate responses of sexual dimorphism to direct and indirect selection. The American Naturalist. 196: 391–405.

12. Cooper, T.F., Rozen D.E. & Lenski, R.E. (2003). Parallel changes in gene expression after 20,000 generations of evolution in *Escherichia coli*. Proceedings of the National Academy of Sciences of the United States of America. 100: 1072–1077.

13. Dean, R. & Mank, J.E. (2016). Tissue specificity and sex-specific regulatory variation permit the evolution of sex-biased gene expression. The American Naturalist. 188: E74–E84.

14. Dobin, A. Davis, C.A., Schlesinger, F., Drenkow, J., Zaleski, C., Jha, S., Batut, P., Chaisson, M. & Gingeras, T.R. (2013). STAR: ultrafast universal RNA-seq aligner. Bioinformatics. 29: 15–21.

15. Dutoit, L., Mugal, C.F., Bolívar, P., Wang, M., Nadachowska-Brzyska, K., Smeds, L., Yazdi, H.P., Gustafsson, L. & Ellegren, H. (2018). Sex-biased gene expression, sexual antagonism and levels of genetic diversity in the collared flycatcher (*Ficedula albicollis*) genome. Molecular Ecology. 27: 3572–3581.

16. Dapper, A.L. & Wade, M.J. (2020). Relaxed selection and the rapid evolution of reproductive genes. Trends in Genetics. 36: 640–649.

17. Edelman G.M. & Gally J.A. (2001). Degeneracy and complexity in biological systems. Proceedings of the National Academy of Sciences of the United States of America. 98: 13763–13768.

18. Ellegren H. & Parsch, J. (2007). The evolution of sex-biased genes and sex-biased gene expression. Nature Reviews Genetics. 8: 689–698.

19. Emlen D.J. (2008). The evolution of animal weapons. *Annual Review of Evology*, Evolution, and Systematics. 39: 387–413.

20. Fairbairn, D.J. (2007). ”Introduction: the enigma of sexual size dimorphism.” In D.J. Fairbairn, W.U. Blanckenhorn & T. Székely (eds), *Sex, Size and Gender Roles: Evolutionary Studies of Sexual Size Dimorphism Sex, Size and Gender Roles*. Oxford Academic.

21. Femenias, M.M., Santos, J.C., Sites, J.W., Avila, L.J. & Morando, M. (2022). ExplorATE: A new pipeline to explore active transposable elements from RNA-seq data. Bioinformatics. 38: 3361–3366.

22. Fischer, E.K., Song, Y., Hughes, K.A., Zhou, W. & Hoke, K.L. (2021). Nonparallel transcriptional divergence during parallel adaptation. Molecular Ecology. 30: 1516–1530.

23. Fisher R. (1930). The genetical theory of natural selection. Oxford: Calendon Press.

24. Fu, L., Niu, B., Zhu, Z., Wu, S. & Li, W. (2012). CD-HIT: Accelerated for clustering the next-generation sequencing data. Bioinformatics. 28: 3150–3152.

25. Ghalambor, C.K., Hoke, K.L., Ruell, E.W., Fischer, E.K., Reznick, D.N. & Hughes, K.A. (2015). Non-adaptive plasticity potentiates rapid adaptive evolution of gene expression in nature. Nature. 525: 372–375.

26. Gosden, T. P., K. L. Shastri, P. Innocenti, & S. F. Chenoweth. (2012). The B-matrix harbors significant and sex-specific constraints on the evolution of multicharacter sexual dimorphism. Evolution. 66: 2106–2116.

27. Grath, S. & Parsch, J. (2016). Sex-biased gene expression. Annual Review of Genetics. 50: 29–44.

28. Griffin, R.M., Dean, R., Grace, J.L., Rydén, P. & Friberg, U. (2013). The shared genome is a pervasive constraint on the evolution of sex-biased gene expression. Molecular Biology and Evolution. 30: 2168–2176.

29. Harrison, P.W., Wright, A.E., Zimmer, F., Dean, R., Montgomery, S.H., Pointer, M.A. & Mank, J.E. (2015). Sexual selection drives evolution and rapid turnover of male gene expression. Proceedings of the National Academy of Sciences of the United States of America. 112: 4393–4398.

30. Hollis, B., Houle, D., Yan, Z., Kawecki, T.J. & Keller, L. (2014). Evolution under monogamy feminizes gene expression in *Drosophila melanogaster*. Nature Communications. 5: 3482.

31. Houle D. & Cheng D. (2021). Predicting the evolution of sexual dimorphism in fene expression. Molecular Biology and Evolution 38: 1847–1859.

32. Huylmans, A.K., Macon, A., Hontoria, F. & Vicoso, B. (2021). Transitions to asexuality and evolution of gene expression in *Artemia* brine shrimp. Proceedings of the Royal Society B: Biological Sciences. 288: 20211720.

33. Hsu, S-K., Belmouaden, C., Nolte, V. & Schlötterer, C. (2020). Parallel gene expression evolution in natural and laboratory evolved populations. Molecular Ecology. 30: 884–894.

34. Hämäläinen, A., Immonen, E., Tarka, M. & Schuett, W. (2018). Evolution of sex-specific pace-of-life syndromes: causes and consequences. Behavioural Biology and Sociobiology. 72: 50.

35. Immonen, E., Snook, R.R. & Ritchie, M.G. (2014). Mating system variation drives rapid evolution of the female transcriptome in *Drosophila pseudoobscura*. Ecology and Evolution. 4: 2186–2201.

36. Immonen, E., Sayadi, A., Bayram, H. & Arnqvist, G. (2017). Mating changes sexually dimorphic gene expression in the seed beetle *Callosobruchus maculatus*. Genome Biology and Evolution. 9: 677–699.

37. Immonen E., Hämäläinen, A., Schuett, W. & Tarka, M. (2018). Evolution of sex-specific pace-of-life syndromes: genetic architecture and physiological mechanisms. Behavioural Ecology and Sociobiology. 72: 60.

38. Immonen, E., Sayadi, A., Stojković, B., Savković, U, Đorđević, M., Liljestrand-Rönn, J., Wiberg, R.A.W. & Arnqvist, G. (2023). Experimental life history evolution results in sex-specific evolution of gene expression in seed beetles. Genome Biology and Evolution. 15: evac177.

39. Ingleby, F.C., Flis, I. & Morrow, E.H. (2015). Sex-biased gene expression and sexual conflict throughout development. Cold Spring Harbour Perspectives in Biology. 7: a017632.

40. Jacobs, A., Carruthers, M., Yurchenko, A., Gordeeva, N.V., Alekseyev, S.S., Hooker, O., Leong, J.S., Minkley, D.R., Rondeau, E.B., Koop, B.F., Adams, C.E. & Elmer, K.R. (2020). Parallelism in eco-morphology and gene expression despite variable evolutionary and genomic backgrounds in a Holarctic fish. PLoS Genetics. 16: e1008658.

41. Jeukens, J., Renaut, S., St-Cyr, J., Nolte, A. W. & Bernatchez, L. (2010). The transcriptomics of sympatric dwarf and normal lake whitefish (Coregonus clupeaformis spp., Salmonidae) divergence as revealed by next-generation sequencing. Molecular Ecology.19, 5389–5403.

42. José Rivas, M., Saura, M., Pérez-Figueroa, A., Panova, M., Johansson, T., André, C., Caballero, A., Rolán-Alvarez, E., Johanneson, K. & Quesada, H. (2018). Population genomics of parallel evolution in gene expression and gene sequence during ecological adaptation. Scientific Reports. 8: 16147.

43. Kaufmann, P., Wolak, M.E., Husby, A. & Immonen, E. (2021). Rapid evolution of sexual size dimorphism facilitated by Y-linked genetic variance. Nature Ecology and Evolution. 5: 1394–1402.

44. Kaufmann, P., Howie, J.M. & Immonen E. (2023a). Sexually antagonistic selection maintains genetic variance when sexual dimorphism evolves. Proceedings of the Royal Sociey B: Biology. 290: 20222484.

45. Kaufmann P., Wiberg, R.A.W., Papachristos, K., Scofield, D.G., Tellgren-Roth, C. & Immonen, E. (2023b). Y-linked copy number polymorphism of target of rapamycin (TOR) is associated with sexual size dimorphism in seed beetles. Molecular Biology and Evolution. 40: msad167.

46. Khila, A., Abouheif, E. & Rowe, L. (2012). Function, developmental genetics, and fithness consequences of a sexually antagonistic trait. Science. 336: 585–589.

47. Kopylova, E., Noé, L. & Touzet, H. (2012). SortMeRNA: Fast and accurate filtering of ribosomal RNAs in metatranscriptomic data. Bioinformatics. 28: 3211–3217.

48. Lande R. (1980). Sexual dimorphism, sexual selection, and adaptation in polygenic characters. Evolution. 34: 292–305.

49. Li, W. & Godzik, A. (2006). Cd-hit: A fast program for clustering and comparing large sets of protein or nucleotide sequences. Bioinformatics. 22: 1658–1659.

50. Mank, J.E. (2017). The transcriptional architecture of phenotypic dimorphism. Nature Ecology and Evolution. 1: 0006.

51. Martin, M. (2011). Cutadapt removes adapter sequences from high-throughput sequencing reads. EMBnet J. 17: 10–12.

52. Martin, S.H., Dasmahapatra, K.K., Nadeau, N.J., Salazar, C., Walters, J.R., Simpson, F., Blaxter, M., Manica, A., Mallet, J. & Jiggins, C.D. (2013). Genome-wide evidence for speciation with gene flow in *Heliconius* butterflies. Genome Research. 23: 1817–1828.

53. McGirr, J.A. & Martin, C.H. (2018). Parallel evolution of gene expression between trophic specialists despite divergent genotypes and morphologies. Evolution Letters. 2: 67–75.

54. Meiklejohn, C.D., Parsch, J., Ranz, J.M. & Hartl, D.L. (2003). Rapid evolution of male-biased gene expression in *Drosophila*, Proceedings of the National Acadaemy of Sciences of the United States of America. 100: 9894–9899.

55. Mishra, P., Rundle, H.D. & Agrawal, A.F. (2024). The evolution of sexual dimorphism in gene expression in response to a manipulation of mate competition. Evolution. 78: 746–757.

56. Nijhout, H.F., Riddiford, L.M., Mirth, C., Shingleton, A.W., Suzuki, Y., Callier, V. (2014). The Developmental Control of Size in Insects. Wiley Interdisciplinary Reviews: Developmental Biology. 3:113–134.

57. Parisi, M., Nuttall, R., Naiman, D., Bouffard, G., Malley, J., Andrews, J., Eastman, S. & Oliver B. (2003). Paucity of genes on the *Drosophila* X chromosome showing male-biased expression. Science. 299: 697–700.

58. Parker, D.J., Bast, J., Jalvingh, K., Dumas, Z., Robinson-Rechavi, M. & Schwander, T. (2019). Sex-biased gene expression is repeatedly masculinised in asexual females. Nature Communications. 10: 4638.

59. Parsch, J. & Ellegren, H. (2013). The evolutionary causes and consequences of sex-biased gene expression. Nature Reviews Genetics. 14: 83–87.

60. Patro, R., Duggal, G., Love, M.I., Irizarry, R.A. & Kingsford, C. (2017). Salmon provides fast and bias-aware quantification of transcript expression. Nature Methods. 14: 417–419.

61. Pennell, T.M., Mank, J.E., Alonzo, S.H. & Hosken, D.J. (2024). On the resolution of sexual conflict over shared traits. Proceedings of the Royal Society B. 291: 20240438.

62. Pennell, T.M. & Morrow, E.H. (2013). Two sexes, one genome: The evolutionary dynamics of intralocus sexual conflict. Ecology and Evolution. 3: 1819–1834.

63. Pointer, M.A., Harrison, P.W., Wright, A.E. & Mank, J.E. (2013). Masculinization of gene expression is associated with exaggeration of male sexual dimorphism. 9: e1003697.

64. Price, P.D., Palmer Droguette, D.H., Taylor, J.A., Kim, D.W., Place, E.S., Rogers, T.F., Mank, J.E., Cooney, C.R., Wright, A.E. (2022). Detecting signatures of selection on gene expression. Nature Ecology and Evolution. 6: 1035–1045.

65. Pruesse, E. Quast, C., Knittel, K. Fuchs, B.M., & Ludwig, W. & Peplies, J. & Glöckner, F.O. (2007). SILVA: A comprehensive online resource for quality checked and aligned ribosomal RNA sequence data compatible with ARB. Nucleic Acids Research. 35: 7188–7196.

66. Quast, C., Pruesse, E., Yilmaz, P., Gerken, J., Schweer, T., Yarza, P., Peplies, J. & Glöckner, F.O. (2013). The SILVA ribosomal RNA gene database project: Improved data processing and web-based tools. Nucleic Acids Research. 41: 590–596.

67. R Core Team (2024). R: A Language and Environment for Statistical Computing. R Foundation for Statistical Computing, Vienna, Austria. https://www.R-project.org

68. Robinson, M.D., McCarthy, D.J. & Smyth, G.K. (2010). edgeR: A Bioconductor package for differential expression analysis of digital gene expression data. Bioinformatics. 26: 139–140.

69. Sayadi, A., Barrio, A.M., Immonen, E., Dainat, J., Berger, D., Tellgren-Roth, C., Nystedt, B. & Arnqvist, G. (2019). The genomics footprint of conflict. Nature Ecology and Evolution. 3: 1725–1730.

70. Scharmann, M., Rebelo, A.G. & Pannell, J.R. (2021). High rates of evolution preceded shifts to sex-biased gene expression in *Leucadendron*, the most sexually dimorphic angiosperms. eLife. 10: e67485.

71. Schlötterer, C. (2023). How predictable is adaptation from standing genetic variation? Experimental evolution in Drosophila highlights the central role of redundancy and linkage disequilibrium. Philosophical Transactions of the Royal Society B. 378: 20220046.

72. Srivastava, A., Malik, L., Sarkar, H., Zakeri, M., Almodaresi, F., Soneson, C., Love, M.I., Kingsford, C. & Patro, R. (2020). Alignment and mapping methodology influence transcript abundance estimation. Genome Biology. 21: 239.

73. Texada, M.J., Koyama, T. & Rewitz, K. (2020). Regulation of Body Size and Growth Control. Genetics. 216: 269–313.

74. Thorhölladottir, D.A.V., Nolte, V. & Schlötterer, C. (2023). Temperature-driven gene expression evolution in natural and laboratory populations highlights the crucial role of correlated fitness effects for polygenic adaptation. Evolution. 77: 2081–2089.

75. Toubiana, W., Dechaud, C., Arbore, R. & Khila, A. (2021). Impact of trait exaggeration on sex-biased gene expression and genome architecture in a water strider. BMC Biology. 19: 89.

76. Tosto, N.M., Beasley, E.R., Wong, B.B.M., Mank, J.E. & Flanagan, S.P. (2023). The roles of sexual selection and sexual conflict in shaping patterns of genome and transcriptome variation. Nature Ecology & Evolution. 7: 981–993.

77. Veltsos, P., Fang, Y., Cossins, A.R., Snook, R.R. & Ritchie, M.G. (2017). Mating system manipulation and the evolution of sex-biased gene expression in *Drosophila*. Nature Communications. 8: 2072.

78. Veltsos, P., Porcelli, D., Fang, Y., Cossins, A.R., Ritchie, M.G. & Snook, R.R. (2022). Experimental sexual selection reveals rapid evolutionary divergence in sex-specific transcriptomes and their interactions following mating. Molecular Ecology. 31: 3374–3388.

79. Wedell, N., Kvarnemo, C., Lessels, C.M. & Tregenza, T. (2006). Sexual conflict and life histories. Animal Behaviour. 71: 999–1011.

80. Wyman, M.J., Stinchcombe, J.R. & Rowe, L. (2013). A multivariate view of the evolution of sexual dimorphism. Journal of Evolutionary Biology. 26: 2070–2080.

81. Wright, A.E., Fumagalli, M., Cooney, C.R., Bloch, N.I., Vieira, F.G., Buechel, S.D., Kolm, N.D. & Mank, J.E. (2018). Male-biased gene expression resolves sexual conflict through the evolution of sex-specific genetic architecture. Evolution Letters. 2: 52–61.

82. Yang, X. & Xu, T. (2011). Molecular mechanisms of size control in development and human diseases. Cell Research. 21: 715–729.

83. Yang, L., Zhang, Z. & He, S. (2016). Both male-biased and female-biased genes evolve faster in fish genomes. Genome Biology and Evolution. 8: 3433–3445.

84. Yeaman, S., Hodgins, K.A., Lotterhos, K.E., Suren, H., Nadeau, S., Degner, J.C., Nurkowski, K.A., Smets, P., Wang, T., Gray, L.K., Liepe, K.J., Hamann, A., Holliday, J.A., Whitlock, M.C., Riesberg, L.H., Aitken, S.N. (2016). Convergent local adaptation to climate change in distantly related conifers. Science. 353: 1431–1433.

85. Zhao, L., Wit, J., Svetec, N. & Begun, D. J. (2015). Parallel gene expression differences between low and high latitude populations of *Drosophila melanogaster* and *D. Simulans*. PLoS Genetics. 11: e1005184.

