## supplementary materials for "Sex-biased gene expression under sexually antagonistic and sex-limited selection"

**Content:**

Tables S1-3, S5-9

Figures S1-9

Supplementary methods

**Table S1.** The distributions of DE transcripts across chromosomes in contrasts. Data are split by whether expression was higher in $SL_m$↓/SA (up-in-$SL_m$↓ and up-in-SA) or $SL_m$↑ (up-in-$SL_m$↑). In each case, the number of transcripts on autosomes, and the X-chromosome are given. The "$Chi^2$" columns give the results of a Chi-squared test using the distribution of all genes across chromosomes as the expected distribution. An asterisk (*) denotes a value larger than expected. The expected proportions are: autosomes: 0.95, X-chromosome: 0.05

| **$SL_m$↓ vs. $SL_m$↑** | | | | |
|---|---|---|---|---|
| | **up-in-$SL_m$↓** | **$Chi^2$** | **up-in-$SL_m$↑** | **$Chi^2$** |
| **Males** | 1,452* : 54 [0.96 : 0.04] | $X^2_1 = 10.5$ $p = 0.001$ | 1,280* : 56 [0.96 : 0.04] | $X^2_1 = 4.32$ $p = 0.04$ |
| **Females** | 854* : 19 [0.98 : 0.02] | $X^2_1 = 18.5$ $p < 0.001$ | 913* : 34 [0.96 : 0.04] | $X^2_1 = 6.57$ $p = 0.01$ |
| **SA vs $SL_m$↑** | | | | |
| | **up-in-SA** | **$Chi^2$** | **up-in-$SL_m$↑** | **$Chi^2$** |
| **Males** | 760* : 19 [0.98 : 0.02] | $X^2_1 = 14.0$ $p < 0.001$ | 751* : 20 [0.98 : 0.02] | $X^2_1 = 12.4$ $p < 0.001$ |
| **Females** | 308* : 6 [0.98 : 0.02] | $X^2_1 = 7.74$ $p = 0.005$ | 334* : 2 [0.99 : 0.01] | $X^2_1 = 15.5$ $p < 0.001$ |

**Table S2.** Counts of differentially expressed transcripts in males and females in the contrasts between selection for larger and smaller males via male-limited ($SL_m$↓ vs. $SL_m$↑) and SA selection (SA vs. $SL_m$↑). Counts are separated by higher relative expression in each line (up-in-$SL_m$↓ and up-in-SA, or up-in-$SL_m$↑). The column "Test of equal proportions" compares the proportion of transcripts that are DE, out of all tested transcripts (N = 19,373), between males and females.

| **$SL_m$↓ vs. $SL_m$↑** | | | |
|---|---|---|---|
| | **up-in-$SL_m$↓** | **up-in-$SL_m$↑** | **Test of equal proportions** |
| *Males* | 1,506 | 1,336 | $X^2_1 = 254.2$, $p < 0.001$ |
| *Females* | 873 | 947 | |
| **SA vs. $SL_m$↑** | | | |
| | **up-in-SA** | **up-in-$SL_m$↑** | **Test of equal proportions** |
| *Males* | 779 | 771 | $X^2_1 = 389.5$, $p < 0.001$ |
| *Females* | 314 | 336 | |

**Table S3.** The number of transcripts that are **unique** $SL_m\downarrow$, **parallel**, or **unique SA** across contrasts for males and females.

| **unique** $SL_m\downarrow$ | | | |
|---|---|---|---|
| | **up-in-**$SL_m\downarrow$ | **up-in-**$SL_m\uparrow$ | **total** |
| **Males** | 1,105 | 971 | 2,076 |
| **Females** | 758 | 811 | 1,569 |
| **parallel** | | | |
| | **up-in-**$SL_m\downarrow$**/ SA** | **up-in-**$SL_m\uparrow$ | **total** |
| **Males** | 396 | 363 | 759 |
| **Females** | 115 | 134 | 249 |
| **unique SA** | | | |
| | **up-in-SA** | **up-in-**$SL_m\uparrow$ | **total** |
| **Males** | 411 | 373 | 784 |
| **Females** | 232 | 258 | 490 |

**Table S4.** GO-analyses, see the separate table.

**Table S5.** The distributions of DE transcripts across sex-bias categories in contrasts. Data are split by whether expression was higher in $SL_m\downarrow$/SA (up-in-$SL_m\downarrow$ and up-in-SA) or $SL_m\uparrow$ (up-in- $SL_m\uparrow$). In each case, the number of transcripts that are female-biased, male-biased, or unbiased are given, the proportions are given in square brackets. The "Chi$^2$" columns give the results of a Chi-squared test using the distribution of all genes across sex-bias categories as the expected distribution. The expected proportions are; female-biased: 0.23, male-biased: 0.24, unbiased: 0.53. An asterisk denotes values larger than expected.

| **$SL_m\downarrow$ vs $SL_m\uparrow$** | | | | |
|---|---|---|---|---|
| | **up-in-$SL_m\downarrow$** | **Chi$^2$** | **up-in-$SL_m\uparrow$** | **Chi$^2$** |
| **Males** | 266 : 605* : 635<br>[0.18 : 0.40 : 0.42] | $X^2_2 = 200.79$<br>$p < 0.001$ | 315 : 362* : 659<br>[0.24 : 0.27 : 0.49] | $X^2_2 = 7.29$<br>$p = 0.03$ |
| **Females** | 443* : 108 : 322<br>[0.51 : 0.12 : 0.37] | $X^2_2 = 396.66$<br>$p < 0.001$ | 95 : 427* : 425<br>[0.10 : 0.45 : 0.45] | $X^2_2 = 242.65$<br>$p < 0.001$ |
| **SA vs $SL_m\uparrow$** | | | | |
| | **up-in-SA** | **Chi$^2$** | **up-in-$SL_m\uparrow$** | **Chi$^2$** |
| **Males** | 146 : 283* : 342<br>[0.19 : 0.37 : 0.44] | $X^2_2 = 62.37$<br>$p < 0.001$ | 166 : 170 : 443<br>[0.21 : 0.22 : 0.57] | $X^2_2 = 5.32$<br>$p = 0.07$ |
| **Females** | 64 : 81 : 191<br>[0.19 : 0.24 : 0.57] | $X^2_2 = 3.03$<br>$p = 0.22$ | 53 : 88 : 173*<br>[0.17 : 0.28 : 0.55] | $X^2 = 6.61$<br>$p = 0.04$ |

**Table S6.** Counts of differentially expressed transcripts in males and females in the contrasts between selection for larger and smaller females via male- and female-limited ($SL_m$↓ vs. $SL_f$↑) and male-limited selection ($SL_m$↓ vs. $SL_m$↑). Counts are separated by higher relative expression in each line (up-in-$SL_m$↓ and up-in-$SL_m$↓, or up-in-$SL_f$↑). The column "Test of equal proportions" compares the proportion of transcripts that are DE, out of all tested transcripts (N = 19,373), between males and females.

| **$SL_m$↓ vs. $SL_f$↑** | | | |
|---|---|---|---|
| | **up-in-$SL_m$↓** | **up-in-$SL_f$↑** | **Test of equal proportions** |
| *Males* | 273 | 377 | $X^2_1 = 0.95$, $p = 0.33$ |
| *Females* | 273 | 413 | |
| **$SL_m$↓ vs. $SL_m$↑** | | | |
| | **up-in-$SL_m$↓** | **up-in-$SL_m$↑** | **Test of equal proportions** |
| *Males* | 1,506 | 1,336 | $X^2_1 = 254.2$, $p < 0.001$ |
| *Females* | 873 | 947 | |

**Table S7.** The number of transcripts that are **unique** $\mathbf{SL_f}$**↑**, **parallel**, or **unique** $\mathbf{SL_m}$**↑** across contrasts for males and females.

| **unique** $\mathbf{SL_f}$**↑** | | | |
|---|---|---|---|
| | **up-in-**$\mathbf{SL_m}$**↓** | **up-in-**$\mathbf{SL_f}$**↑** | **total** |
| **Males** | 126 | 157 | 283 |
| **Females** | 154 | 202 | 356 |
| **parallel** | | | |
| | **up-in-**$\mathbf{SL_m}$**↓** | **up-in-**$\mathbf{SL_f}$**↑/**$\mathbf{SL_m}$**↑** | **total** |
| **Males** | 147 | 219 | 366 |
| **Females** | 118 | 210 | 228 |
| **unique** $\mathbf{SL_m}$**↑** | | | |
| | **up-in-**$\mathbf{SL_m}$**↓** | **up-in-**$\mathbf{SL_m}$**↑** | **total** |
| **Males** | 1,358 | 1,117 | 2,475 |
| **Females** | 754 | 735 | 1,789 |

**Table S8.** The distributions of DE transcripts across sex-bias categories in contrasts. Data are split by whether expression was higher in $SL_m$↑/$SL_f$↑ (up-in-$SL_m$↑ and up-in-$SL_f$↑) or $SL_m$↓ (up-in- $SL_m$↓). In each case, the number of transcripts that are female-biased, male-biased, or unbiased are given, the proportions are given in square brackets. The "$Chi^2$" columns give the results of a Chi-squared test using the distribution of all genes across sex-bias categories as the expected distribution. The expected proportions are; female-biased: 0.23, male-biased: 0.24, unbiased:0.53. An asterisk denotes values larger than expected.

| **$SL_m$↓ vs $SL_f$↑** | | | | |
|---|---|---|---|---|
| | **up-in-$SL_m$↓** | **$Chi^2$** | **up-in-$SL_f$↑** | **$Chi^2$** |
| **Males** | 37 : 77* : 159*<br>[0.14 : 0.28 : 0.58] | $X^2_2 = 13.13$<br>$p < 0.001$ | 63 : 127* : 187<br>[0.17 : 0.34 : 0.50] | $X^2_2 = 19.76$<br>$p < 0.001$ |
| **Females** | 87* : 41 : 145<br>[0.32 : 0.15 : 0.53] | $X^2_2 = 20.09$<br>$p < 0.001$ | 51 : 157* : 205<br>[0.12 : 0.38 : 0.50] | $X^2_2 = 51.21$<br>$p < 0.001$ |
| **$SL_m$↓ vs $SL_m$↑** | | | | |
| | **up-in-$SL_m$↓** | **$Chi^2$** | **up-in-$SL_m$↑** | **$Chi^2$** |
| **Males** | 266 : 605* : 635<br>[0.18 : 0.40 : 0.42] | $X^2_2 = 200.79$<br>$p < 0.001$ | 315 : 362* : 659<br>[0.24 : 0.27 : 0.49] | $X^2_2 = 7.29$<br>$p < 0.03$ |
| **Females** | 443* : 108 : 322<br>[0.51 : 0.12 : 0.37] | $X^2_2 = 396.66$<br>$p < 0.001$ | 95 : 427* : 425<br>[0.10 : 0.45 : 0.45] | $X^2_2 = 242.65$<br>$p < 0.001$ |

**Table S9.** Associations with sex-bias among differentially expressed transcripts in females (see main text for details). Counts with asterisks indicate a significantly greater proportion than expected based on genome-wide proportions of sex-biased transcripts (shown in brackets in the column headers). Proportions out of row totals are given in brackets. Distributions are significantly different across rows ($X^2_6$ = 131, p < 0.001).

| | **Female-biased (0.23)** | **Male-biased (0.24)** | **Unbiased (0.53)** | **total** |
|---|---|---|---|---|
| *not-DE* | 3,649 (0.23) | 3,597 (0.23) | 8,405 (0.54) | 15,754 |
| unique $SL_f$↑ | 68 (0.19) | 97 (0.27) | 191 (0.54) | 356 |
| *parallel* | 70 (0.21) | 100* (0.30) | 158 (0.48) | 328 |
| *unique* $Sl_m$↑ | 468 (0.31) | 434* (0.29) | 588 (0.39) | 1,490 |

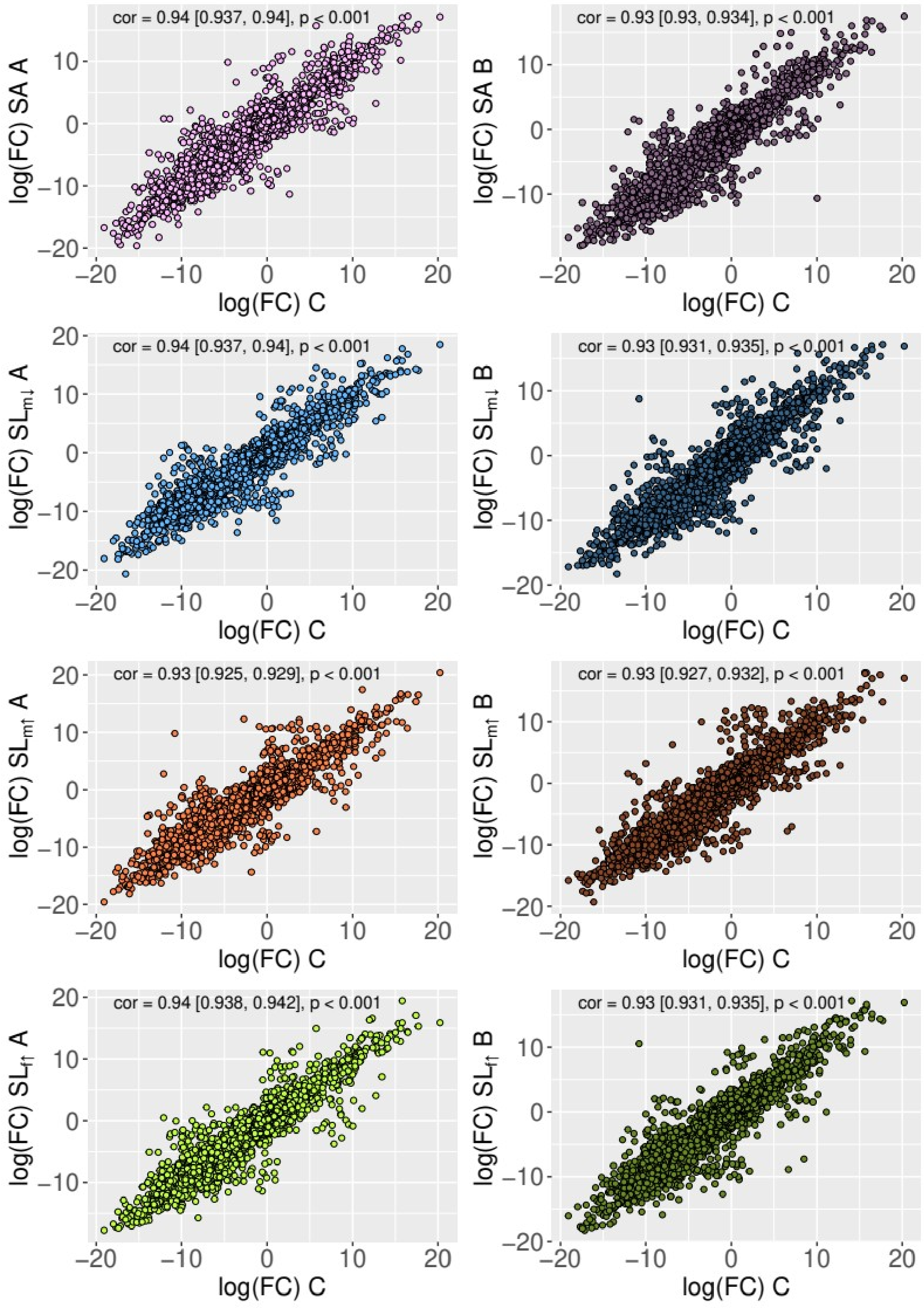


**Figure S1.** Correlation in the sex-difference in expression between the C line and each selection treatment. Numbers inset at the top of each panel give the results of a Pearson's correlation test.

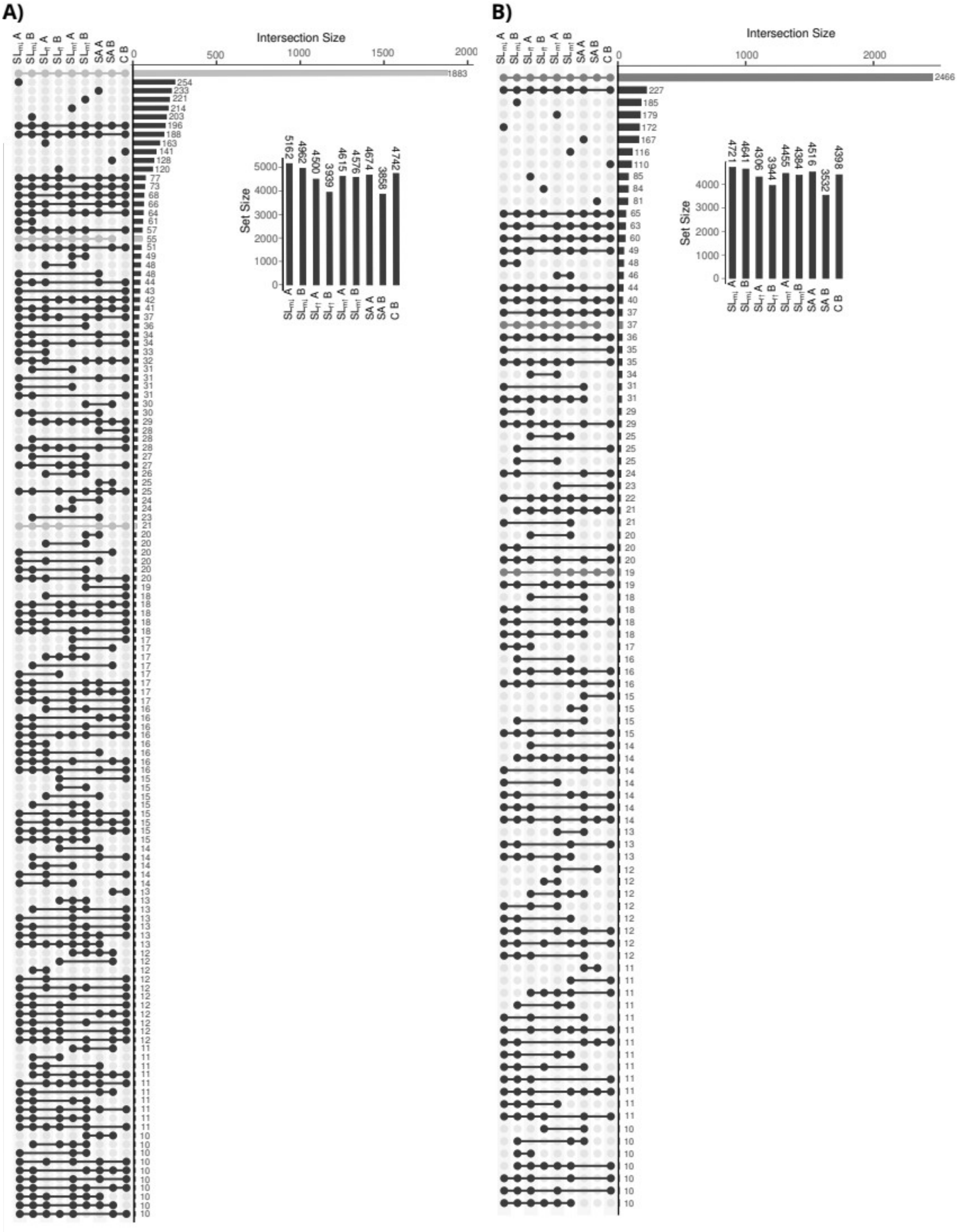


**Figure S2.** UpSet plot of the overlap in sex-biased transcripts across selection lines. Highlighted are intersections of transcripts in both replicates of L1 or SA selection lines and at least 2 other selection lines. **A)** male-biased transcripts and **B)** female-biased transcripts.

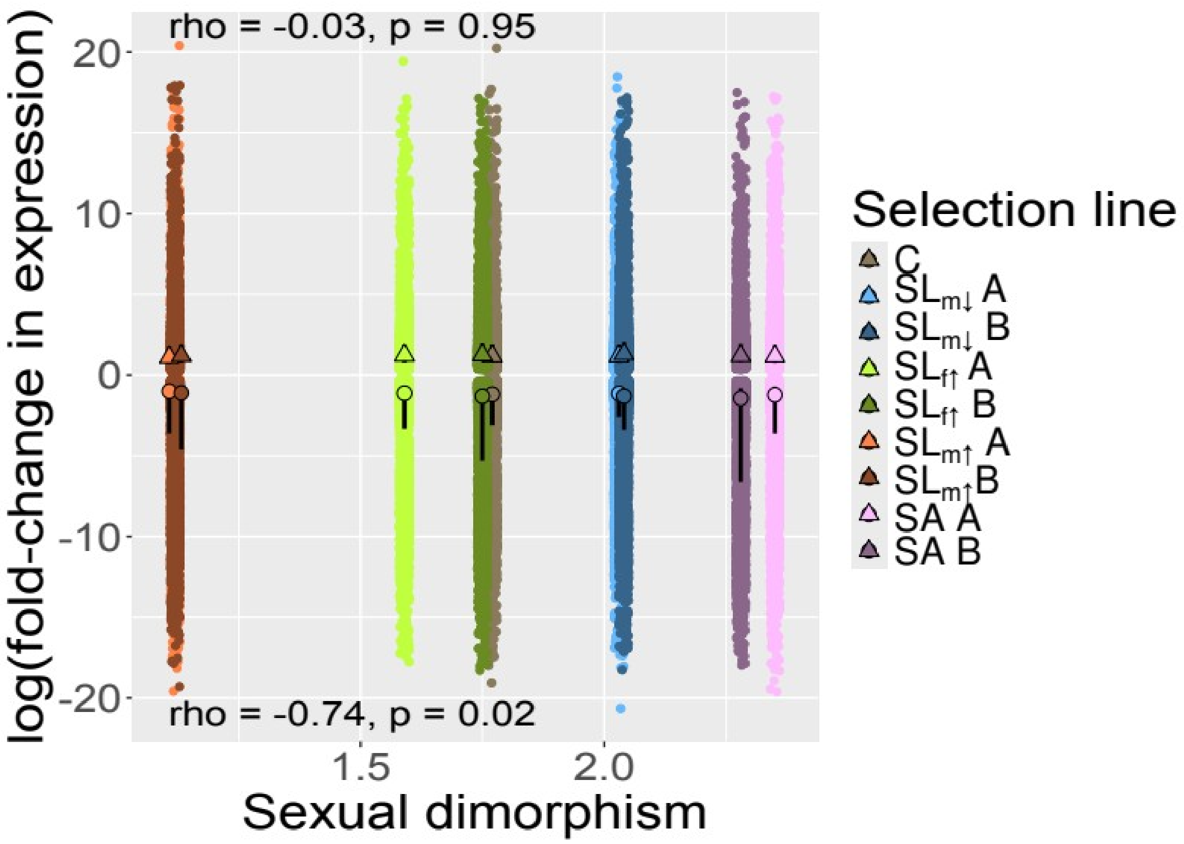


**Figure S3.** log fold-change (FC) in expression between males and females for transcripts with significant sex-bias in each selection treatment as a function of sexual size dimorphism. Red triangles and circles give the median logFC for male- and female- biased transcripts respectively. Inset text at the top and bottom of the figure give results for male- and female-biased transcripts, respectively, of a Spearman rank correlation test of median logFC values on sexual dimorphism.

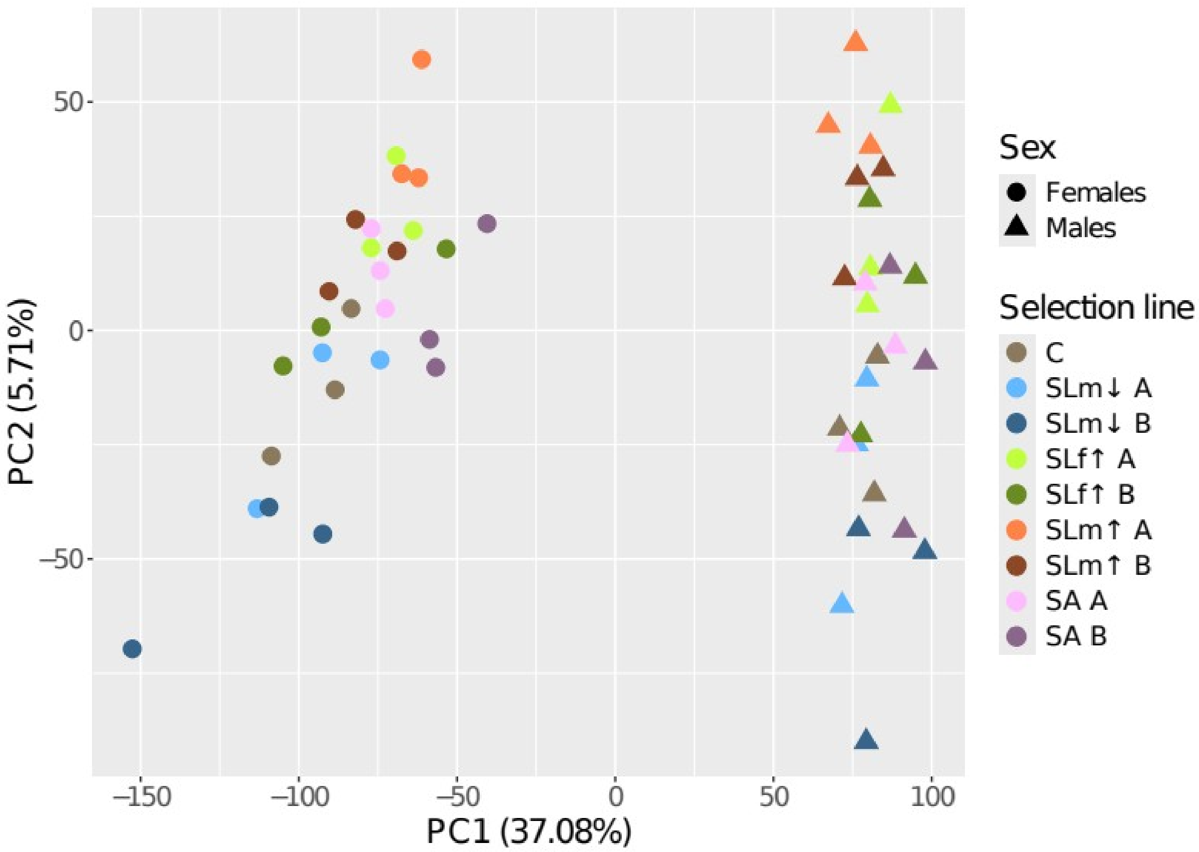


**Figure S4.** Biplot of the first and second principle components from a PCA of all expression data across samples. Points are separated by males and females (shape) and selection line (colour).

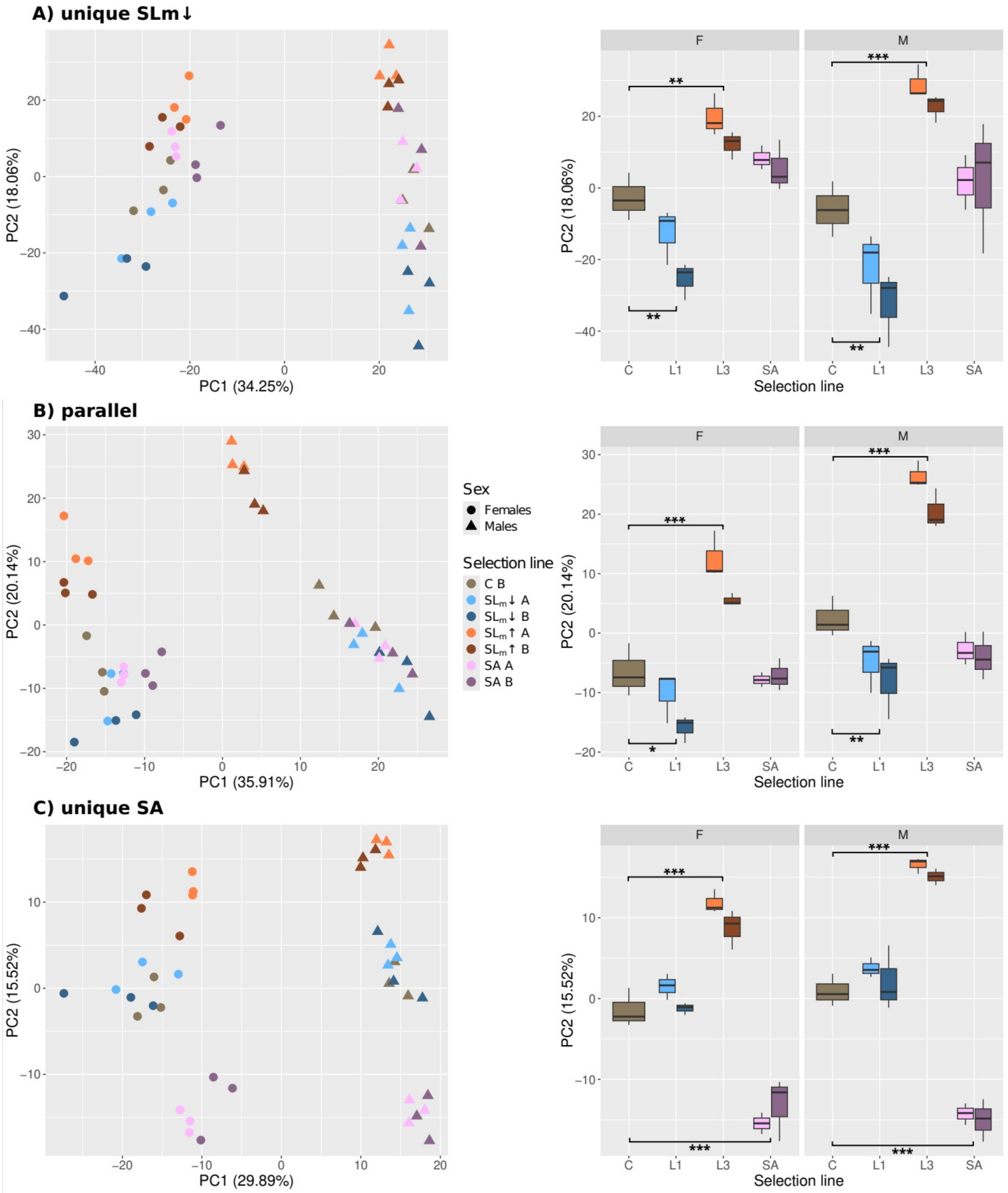


**Figure S5.** The left column of plots shows PCs 1 and 2 of a PCA of expression across **A)** *unique* $SL_m$↓ **B)** *parallel*, and **C)** *unique* SA transcripts. The right column shows the distribution of values of PC2 as a function of selection line. Contrasts marked with a bar and asterisks are statistically significant in a t-test at * $p < 0.05$, ** $p < 0.01$, *** $p < 0.001$.

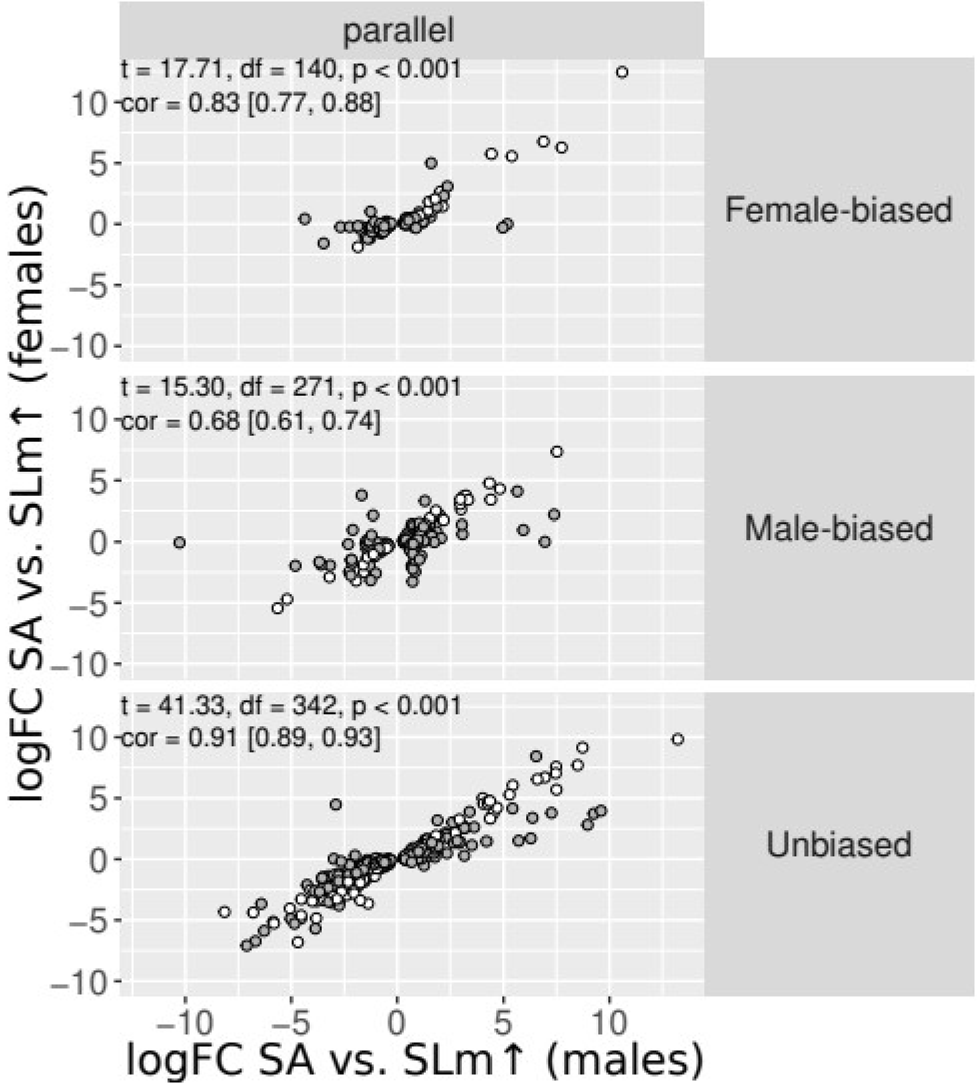


**Figure S6.** The logFC values for males and females of the parallel transcript class in the SA vs. $SL_m$↑. Data are split by sex-bias status. Points are coloured by whether the sexes show the same expression response (white) or not (grey). Inset text gives results of Pearson's correlation tests as well as correlation coefficients for each panel.

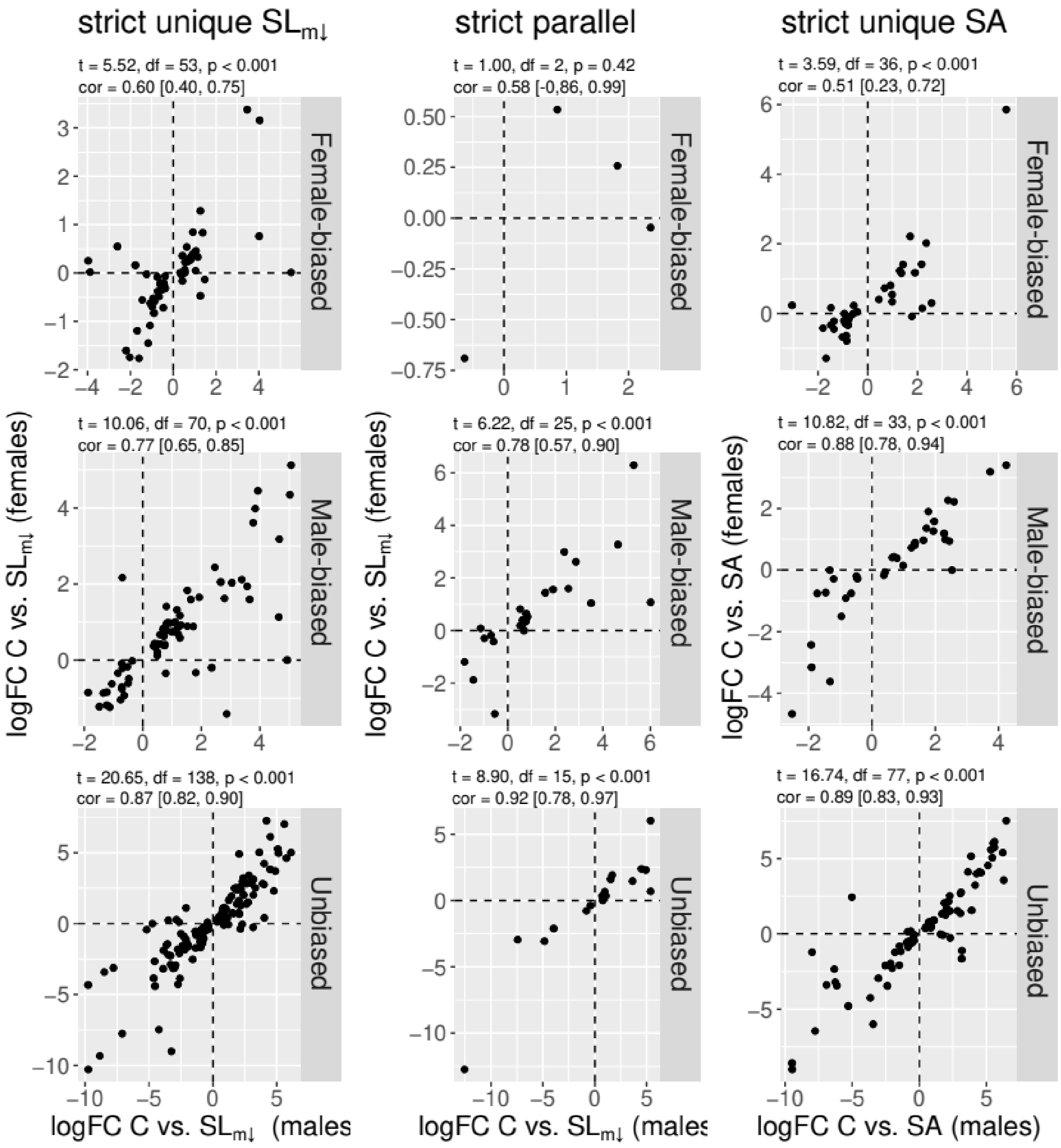


**Figure S7.** LogFC values in C vs. SL$_m$↓ and C vs. SA contrasts for each transcript class and sex-bias status. Across panels, positive values indicate higher expression in the C line. Inset text above each panel shows the results of a Pearson correlation test of male on female logFC values for transcripts within each panel. The 95% CI is given for each correlation coefficient in squared brackets. For one panel (female-biased, strict parallel), inset text is shown in red, here the correlation test was performed while excluding the highlighted point (in red).

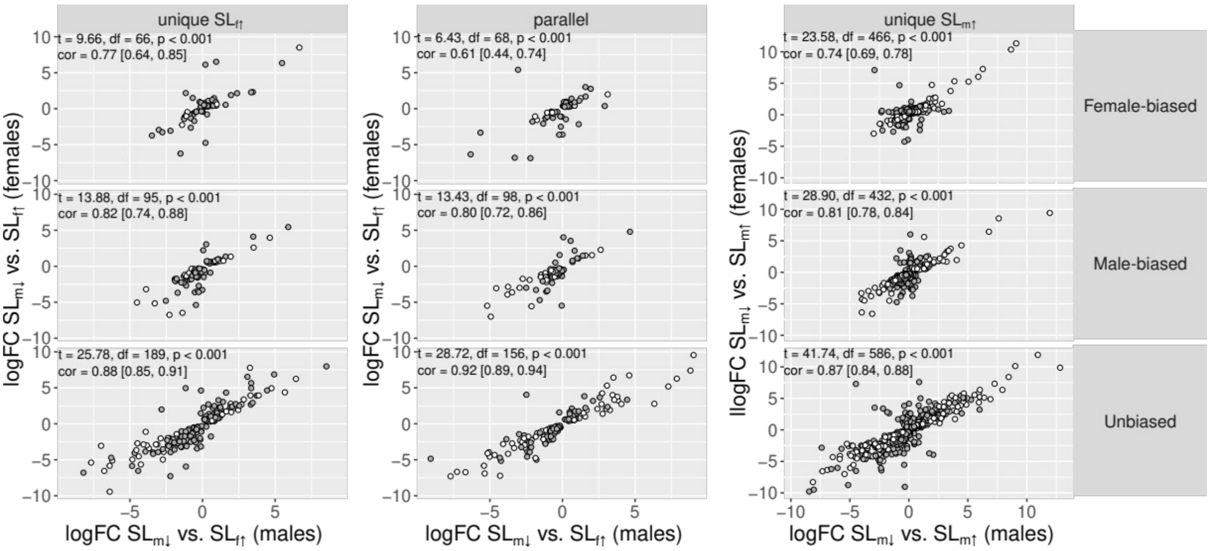


**Figure S8.** logFC expression between $SL_m\downarrow$ and $SL_f\uparrow$, or $SL_m\downarrow$ and $SL_m\uparrow$, for males and females. Data are split by whether males show a unique response in the $SL_m\downarrow$ vs. $SL_f\uparrow$ contrast, a unique response in the $SL_m\downarrow$ vs. $SL_m\uparrow$ contrast, or a parallel response in both, and by sex-bias status. Points are coloured by whether both of the sexes show the same significant expression difference (white) or only one of them (grey). Inset text gives results of Pearson's correlation tests as well as correlation coefficients for each panel. Note that the axis labels are different for the final column of panels. The logFC values for males and females of the parallel transcript class in the $SL_m\downarrow$ vs. $SL_m\uparrow$ contrast is not shown.

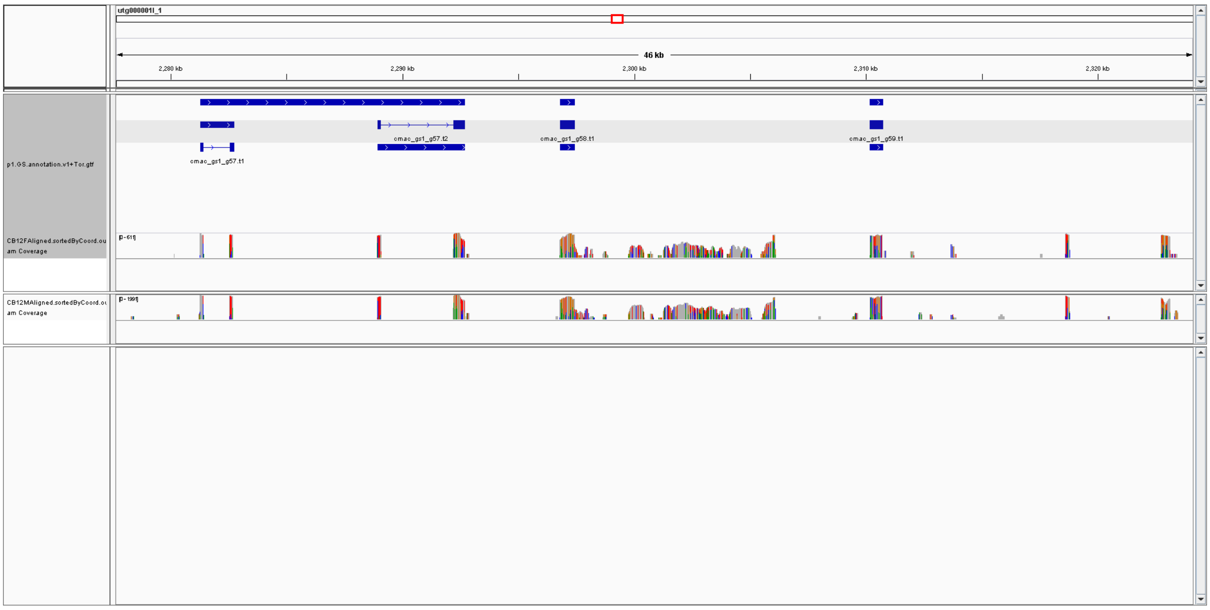


**Figure S9.** Representative screen-shot from the IGV. Shown are annotated gene models with intron/exon boundaries in blue boxes. Coverage tracks from two RNA-Seq samples are also shown and indicate RNA-Seq reads mapping to genomic regions not annotated with genes.

### Supplementary Methods

*Read processing, mapping, overall expression patterns*

Mapping rates were unexpectedly low at 46%. SortMeRNA (v. 4.3.3; Kopylova et al. 2012) identified between 7% and 14% as coming from rRNA sequences. Mapping rates were negatively correlated with the number of reads, but only for males. We mapped the RNA-seq reads from adult samples of the control (C) lines to the genome assembly. This resulted in much higher mapping rates (76%). Only 0.5% of reads were excluded due to mapping to too many loci. Meanwhile, 18% of reads were excluded due to too many mismatches between the reads and the reference genome. Additionally, ~3% of reads arise from active repeat elements in the genome. Manual inspection of coverage variation throughout the genome in IGV (v.2.15.2; Robinson et al. 2011) indicates that many reads are mapped to intergenic regions (figure S9). In many cases coverage blocks indicate the presence of unannotated exons. Such reads do not map to the transcriptome and thereby make up the majority of the difference between the mapping rates of 40% to the transcriptome and 70% to the genome.

Thus, the relatively low mapping rates can be explained by a combination of incomplete gene annotations, excessive mismatches between reads and the reference genome, as well as active TEs that contribute mRNA fragments to the RNA samples. In sum, expression estimates are based on a mean 24.4 million reads across samples.

In principle component analyses (PCA) of expression across all transcripts and samples, PC1 separates very clearly the samples by sex (figure S5). Moreover, there is also separation of samples from different selection lines along PC2 (figure S5).

### References

Kopylova, E., Noé, L. & Touzet, H. (2012). SortMeRNA: Fast and accurate filtering of ribosomal RNAs in metatranscriptomic data. *Bioinformatics*. **28**: 3211-3217.

Robinson, J.T., Thorvaldsdóttir, H., Winckler, W., Guttman, M., Lander, E.S., Getz, G. & Mesirov, J.P. (2011). Integrative genomics viewer. Nature Biotechnology. 29: 24-26.
